# Overlapping yet Distinct Functions of VMP1 and TMEM41B in Modulating Hepatic Lipoprotein Secretion and Autophagy

**DOI:** 10.1101/2025.04.07.647617

**Authors:** Allen Chen, Khue Nguyen, Xiaoxiao Jiang, Xiaochen Yu, Yan Xie, Wanqing Liu, Nicholas O. Davidson, Wen-Xing Ding, Hong-Min Ni

## Abstract

**Background:** Transmembrane protein 41B (TMEM41B) and vacuolar membrane protein 1 (VMP1) are endoplasmic reticulum (ER) scramblases that shuttle phospholipids between the inner and outer leaflets of the ER membrane. Both TMEM41B and VMP1 also play critical roles in regulating hepatic lipoprotein secretion and autophagy. Despite these similarities, whether TMEM41B and VMP1 exhibit different roles in very low-density lipoprotein (VLDL) secretion and autophagy in the pathogenesis of metabolic-associated steatotic liver disease (MASLD) remains unclear.

**Methods:** We created liver- and hepatocyte-specific single knockout (KO) and double knockout (DKO) mice for *Tmem41b* and *Vmp1*, as well as overexpression knock-in (KI) mice with hepatic overexpression of TMEM41B, *Tmem41b* KO/*Vmp1* KI, and *Vmp1* KO/*Tmem41b* KI. We conducted lipidomic, metabolomic, biochemical, and functional studies in these mice, fed either a chow diet or a MASLD diet.

**Results:** TMEM41B protein levels were decreased in the livers of human subjects with MASLD. The loss of hepatic *Tmem41b* impaired VLDL secretion, resulting in steatosis, inflammation, and fibrosis. *Vmp1* KO mice exhibited similar phenotypes to DKO mice, displaying a more severe defect in VLDL secretion and greater liver injury than *Tmem41b* KO mice. Lipidomic analysis revealed decreased levels of phosphatidylcholine and phosphatidylethanolamine, along with increased neutral lipids in both *Tmem41b* KO and *Vmp1* KO mice; however, these changes were generally more pronounced in *Vmp1* KO mice. VMP1 and TMEM41B localized at the mitochondrial-associated membrane (MAM), and a reduction in mitochondria-ER contact was observed in hepatocytes deficient in either VMP1 or TMEM41B. Ultrastructural electron microscopy analysis showed increased accumulation of “lipid droplet” in the ER membrane bilayer and ER lumen in both *Vmp1* KO and *Tmem41b* KO hepatocytes, with greater ER luminal “lipid droplet” accumulation in *Tmem41b* KO hepatocytes. The loss of hepatic *Vmp1* or *Tmem41b* led to elevated levels of LC3-II and p62, with significantly higher levels of both markers in *Vmp1* KO and DKO mouse livers compared to *Tmem41b* KO mouse livers. Restoring *Vmp1* in *Tmem41b* KO mice partially improved defective VLDL secretion; however, high expression levels of VMP1 did not correct the hepatic autophagy defect. In contrast, restoring *Tmem41b* in *Vmp1* KO mice dose-dependently enhanced both defective VLDL secretion and autophagy. Importantly, overexpression of hepatic TMEM41B mitigated diet-induced MASLD in mice.

**Conclusion:** The loss of hepatic *Vmp1* or *Tmem41b* decreases hepatic MAM and phospholipid content, leading to decreased VLDL secretion and promoting MASLD. While VMP1 and TMEM41B have overlapping functions, VMP1 appears to play a more critical role in regulating VLDL secretion and autophagy in mouse livers than TMEM41B.

## Introduction

Metabolic dysfunction-associated steatotic liver disease (MASLD) has become the most common chronic liver disease globally, affecting an estimated 24% of the population and having limited FDA-approved pharmacotherapies [1, 2]. Left untreated, MASLD can progress to metabolic dysfunction-associated steatohepatitis (MASH), which is defined by hepatocyte death and inflammation in the setting of steatosis. Unaddressed, MASH can precipitate the development of end-stage liver diseases, including cirrhosis and hepatocellular carcinoma [3, 4].

Research into MASLD/MASH treatment commonly focuses on upregulating hepatic lipid catabolism or blocking absorption of lipids from the gut, thus decreasing overall systemic lipid levels [2, 5]. Resmetirom is currently the only FDA approved drug for treating MASH. It is a liver-targeted selective thyroid hormone receptor β agonist. Resmetirom works by promoting the oxidation of fatty acids in the liver, while also inhibiting *de novo* lipogenesis and the production of very low-density lipoproteins (VLDL). This leads to improved lipid metabolism and reduced lipotoxicity in the liver [6–8]. Therefore, targeting hepatic lipid metabolism, including lipid synthesis (de novo), breakdown (beta oxidation), and secretion (VLDL), could potentially improve liver pathology in patients with MASLD/MASH.

VLDL secretion is a complex process. It begins in the endoplasmic reticulum (ER) membrane with the synthesis of apolipoprotein B-100 (apoB-100), the primary structural component of VLDL. Following the synthesis of apoB-100, lipids such as triglycerides and cholesterol esters are incorporated into the apoB-100 molecule within the ER lumen, a process facilitated by the microsomal triglyceride transfer protein (MTTP). After this lipid incorporation, VLDL is packaged into specialized transport vesicles coated with COPII proteins and transported to the Golgi apparatus for further maturation before being secreted into the circulation [9]. Importantly, genetic variants such as PNPLA3 and TM6SF2 are associated with MASLD and both regulate the production and secretion of VLDL, highlighting the significance of VLDL secretion in MASLD [9, 10].

Vacuolar membrane protein 1 (VMP1) and Transmembrane Protein 41B (TMEM41B) are transmembrane proteins located in the ER that function as phospholipid scramblases [11, 12]. They play crucial roles in regulating the formation of autophagosomes and lipid droplets (LDs) [12–15]. Recent research, including our own, has shown that VMP1 and TMEM41B are two novel regulators of hepatic VLDL secretion. Notably, both proteins are found to be decreased in the livers of mice with diet-induced MASH and in human patients with MASH [11, 16]. Mice with single knockouts of *Vmp1* and *Tmem41b* exhibit impaired VLDL secretion and rapidly develop hepatic steatosis, even when fed a standard chow diet [11, 16]. However, it remains unclear whether VMP1 and TMEM41B operate through similar mechanisms or serve distinct roles in VLDL secretion and the development of MASLD. Additionally, it remains uncertain whether VMP1 and TMEM41B can compensate for each other’s functions or if they operate independently or in coordination with one another in their roles regulating VLDL secretion and autophagy. In this study, we investigate the expression of TMEM41B in the livers of subjects with MASH and further explore the role of TMEM41B in relation to VMP1 regarding lipid homeostasis, autophagy, and VLDL secretion in mouse liver. Our findings offer new insights into the molecular relationship between VMP1 and TMEM41B, highlighting their redundant yet distinct roles in VLDL secretion and autophagy as well as hepatic metabolism. Furthermore, we suggest that VMP1 and TMEM41B could be potential targets for treating MASLD and MASH.

## Materials and Methods

### Animals

The *Vmp1^flox^* and *Vmp1* conditional knockin (KI) mice were described previously [14]. The *Tmem41b^flox^* and *Tmem41b* conditional KI mice were generated in collaboration with Cyagen (Santa Clara, CA). Briefly, to generate *Tmem41b^flox^*mice, ES cells containing *Tmem41b* exons 3-5 cassette which were flanked by two loxP sequences. Cre-mediated depletion of exons 3-5 leads to a frameshift, resulting in a small truncated peptide. To generate *Tmem41b* conditional KI mice, ES cells containing an insertion of *Tmem41b-3×Flag* cassette which contains a polyA tail flanked by two *loxP* sequences between CAG promoter and *Tmem41b-3×Flag* under *Rosa26* loci were injected into C57BL/6N blastocysts to obtain chimeric mice. The chimeric mice were crossed with C57BL/6N to obtain mutant mice. Cre-mediated depletion of the polyA tail leads to expression of TMEM41b-FLAG. *Tmem41b^flox,^ Vmp1^flox^* mice were generated by crossing *Tmem41b^flox^* with *Vmp1^flox^*mice. *Tmem41b* ^flox^, *Vmp1*^KI^ mice were generated by crossing *Tmem41b*^flox^ mice with *Vmp1*^KI^ mice. *Vmp1^flox^*, *Tmem41b^KI^* mice were generated by crossing *Vmp1^flox^* mice with *Tmem41b^KI^* mice. To generate liver-specific KO/KI mice, the mutant mice were crossed with albumin-Cre mice. To generate hepatocyte-specific *Tmem41b* (H-*Tmem41b)* KO mice, *Tmem41b^flox^* were injected intravenously with adeno-associated virus 8 (AAV8)-thyroxine binding globulin promoter (TBG)-null or AAV8-TBG-cre (1×10^11^ GC/mouse). All mice were fed with a chow diet unless otherwise indicated. To generate a model of diet-induced MASH, mice were fed a CDAHFD (choline-deficient, amino acid-defined high-fat diet (45% fat) containing 0.1% methionine) for 6 weeks. Mice were specific pathogen-free and maintained in a barrier rodent facility under standard experimental conditions. The Institutional Animal Care and Use Committee of the University of Kansas Medical Center approved all procedures.

### Reagents and antibodies

Antibodies and reagents used in this study are listed in the supplementary Table 1.

### Hepatic triglyceride (TG) and cholesterol analysis

Hepatic lipid extraction was performed as described previously[17]. Briefly, frozen liver tissues (20-50 mg) were homogenized and followed by lipid extraction using a chloroform-methanol mixture. The resulting lipid extracts were analyzed for TG and cholesterol content using the GPO-Triglyceride Reagent Set and Cholesterol liquid reagent (catalog #T7532 and #C7510, Pointe Scientific, Canton, MI), following the manufacturer’s protocols.

### *In vivo* VLDL secretion assay

*In vivo* VLDL secretion assay was performed as previously described[18]. Mice were subjected to a 4-hour fast prior to the intraperitoneal administration of Pluronic™ F-127 at a dosage of 10 mg/kg. Pluronic™ F-127 inhibits both the lipolysis and the tissue uptake of lipoproteins in mice. Blood samples were collected at baseline (pre-injection) and subsequently at hourly intervals for a duration of 3 hours post-injection. TG concentrations were quantified using the colorimetric assay described above.

### Blood Biochemistry analysis

Serum TG and cholesterol levels were quantified using the colorimetric assays outlined above. Serum alanine aminotransferase (ALT) activity was assessed using commercially available kit (catalog #A7526, Pointe Scientific) in accordance with the manufacturer’s instructions.

### Histology and immunohistochemistry analysis

Livers were preserved in 10% neutral buffered formalin and subsequently embedded in paraffin. Sections of 5 μm thickness from the paraffin blocks were stained using hematoxylin and eosin (H&E) for histopathological assessment. For immunohistochemical analysis, F4/80 was utilized to identify macrophages, while Sirius Red staining was employed to visualize collagen deposition.

### Oil Red O staining

Oil Red O staining was performed using liver cryo-sections as described previously[19]. Livers were preserved in 4% paraformaldehyde overnight at 4°C, then infiltrated with 20% sucrose overnight at the same temperature, and embedded in Tissue-Tek OCT Compound. Sections of 6 μm thickness were stained with Oil Red O in 60% isopropanol for 15 minutes at 37°C. This was followed by a wash with 60% isopropanol and three washes with water, then stained with hematoxylin, concluding with a final wash with water.

### Electron microscopy analysis

Electron microscopy was performed as described previously[20]. Breifly, livers were perfused with 2.5% glutaraldehyde in a 0.1M sodium cacodylate buffer and subsequently sectioned into small fragments. Ultra-thin sections were then stained with uranyl acetate and lead citrate. Imaging was performed using a JEM 1016CX electron microscope (JEOL) to analyze the ultrastructural details.

### Immunoblot analysis

Liver proteins were extracted using RIPA buffer, containing 1% NP40, 0.5% sodium deoxycholate, and 0.1% sodium dodecyl sulfate (SDS) in phosphate-buffered saline (PBS). A total of 30 μg of protein was separated on an SDS-PAGE gel, followed by transfer onto a PVDF membrane. The membranes were subsequently incubated with appropriate primary antibodies, followed by corresponding secondary antibodies, and detection was performed using SuperSignal Plus chemiluminescent substrate (Thermo Fisher Scientific). Densitometric analysis of the bands was conducted using ImageJ or Un-Scan-It software, with normalization to β-actin or GAPDH. All densitometry data are reported as mean ± SEM.

### Quantitative real-time polymerase chain reaction

RNA was extracted from murine hepatic tissue using Trizol reagent (Thermo Fisher Scientific) and subsequently reverse transcribed into complementary DNA (cDNA) utilizing RevertAid H minus reverse transcriptase (Thermo Fisher Scientific) [21]. Quantitative real-time PCR was conducted on a Bio-Rad CFX384™ detection system employing SYBR® Green mix (Bimake, Houston, TX). Expression of *Acta2, Adgre, Cd68, Col1a1, Ctgf, Il1b, Il10, Tgfb1, and Trem2* was quantified using qRT-PCR analysis and *Actb* was used as an internal control. The fold change of mRNA was expressed as 2^−ΔΔCt^.

### Lipidomics and metabolomics analysis

Lipidomic and metabolomic profiling of mouse liver tissues was conducted at Metabolon, Inc. (Durham, NC) as described previously [14, 22]. Briefly, for lipidomic analysis, lipids were extracted from samples in a methanol:dichloromethane mixture in the presence of internal standards. After concentration under nitrogen, the lipid extracts were reconstituted in 0.25 mL of 10 mM ammonium acetate dichloromethane:methanol (50:50). These extracts were subsequently transferred to inserts and loaded into vials for infusion mass spectrometry (MS) analysis, utilizing a Shimadzu LC equipped with nano PEEK tubing and the Sciex SelexION-5500 QTRAP. The analysis was conducted in both positive and negative ionization modes. The 5500 QTRAP operated in Multiple Reaction Monitoring (MRM) mode, with over 1,100 monitored reactions. Quantification of individual lipid species was achieved by calculating the peak area ratios of the target lipids to their corresponding internal standards, which were then normalized to the known concentrations of the internal standards added during extraction. Lipid class concentrations were derived as the aggregate of all molecular species within each class, while fatty acid profiles were delineated by determining the constituency of each class based on individual fatty acids. Statistical evaluations were conducted using ArrayStudio, R (http://cran.r-project.org/), or JMP, with significance assessed via one-way ANOVA. The false discovery rate (FDR) was estimated utilizing q-values to adjust for multiple comparisons.

Metabolomics analysis was performed by Metabolon as described previously [22]. Following log transformation and imputation of missing values, if any, with the minimum observed value for each compound, Welch’s two-sample t-test was used to identify biochemicals that differed significantly between experimental groups. A summary of the numbers of biochemicals that achieved statistical significance (p≤0.05), as well as those approaching significance (0.05<p<0.10) is shown.

### Subcellular fractionation

Subcellular fractionation was performed using an adapted, previously described protocol [23]. Mouse livers from 1-month-old LWT, *Tmem41b*^LKO^ and *Vmp1*^LKO^ mice were homogenized in a mannitol-sucrose-BSA-EGTA-Tris buffer using a Potter-Elvehjem tissue homogenizer. Mitochondria were separated from cytosolic fractions through a series of 9,000g-10,000g centrifugations. Following this, 95,000g-100,000g centrifugations in 30% vol/vol Percoll solution was performed to separate MAM from mitochondria. ER was isolated from cytosolic fractions after subjecting the supernatant from a 20,000g centrifugation to a 100,000g centrifugation then collecting pellet. Subcellular fractions were subsequently performed using immunoblot analysis.

### Statistical analysis

Data were analyzed in R or with GraphPad. All experimental data are expressed as mean ± SEM and subjected to unpaired Student’s *t test* (2 group comparisons) or one-way ANOVA with post-hoc Turkey test (multigroup comparisons).

## Results

### Hepatic deletion of TMEM41B impairs VLDL secretion and leads to rapid development of microvesicular Steatosis

To investigate the physiological functions of TMEM41B in mice, 8-week-old *Tmem41b^flox^* mice were injected with a single dose of AAV8-TBG-Cre (hepatocyte-specific *Tmem41b* KO, *Tmem41b*^HKO^) or AAV8-TBG-Null (hepatocyte wild-type (HWT)) for 2 and 4 weeks. Compared to HWT mice, *Tmem41b*^HKO^ mice had enlarged and pale livers with significant increases in liver-body weight ratio (**Fig. 1A**). Hematoxylin and Eosin (H&E) staining revealed pan-hepatic microvesicular steatosis in *Tmem41b*^HKO^ mice (**Fig. 1B**). Hepatic triglyceride (TG) and cholesterol were significantly elevated in *Tmem41b*^HKO^ mice compared with HWT mice (**Fig. 1C**). Interestingly, serum TG and cholesterol levels were significantly decreased post-AAV injection (**Fig. 1D**). Therefore, we subsequently performed lipoprotein secretion analysis in *Tmem41b*^HKO^ and HWT mice. Compared with HWT mice, serum TG was markedly decreased in *Tmem41b*^HKO^ mice in a time-dependent manner with Pluronic™ F-127 administration to inhibit lipolysis and tissue uptake of TG-rich lipoproteins. TG secretion rate was reduced by 81.55% in *Tmem*41b^HKO^ mice (**Fig. 1E**). Immunoblot analysis revealed selectively decreased serum apolipoprotein (APO)B100 in *Tmem41b*^HKO^ mice with preserved secretion of APOB48 (**Fig. 1F**). Fast protein liquid chromatography (FPLC) analysis demonstrated substantially decreased VLDL-TG levels in *Tmem41b*^HKO^ mice without affecting peak distribution (**Fig. 1G**), confirming impaired VLDL secretion in *Tmem41b*^HKO^ mice.

**Fig. 1.**
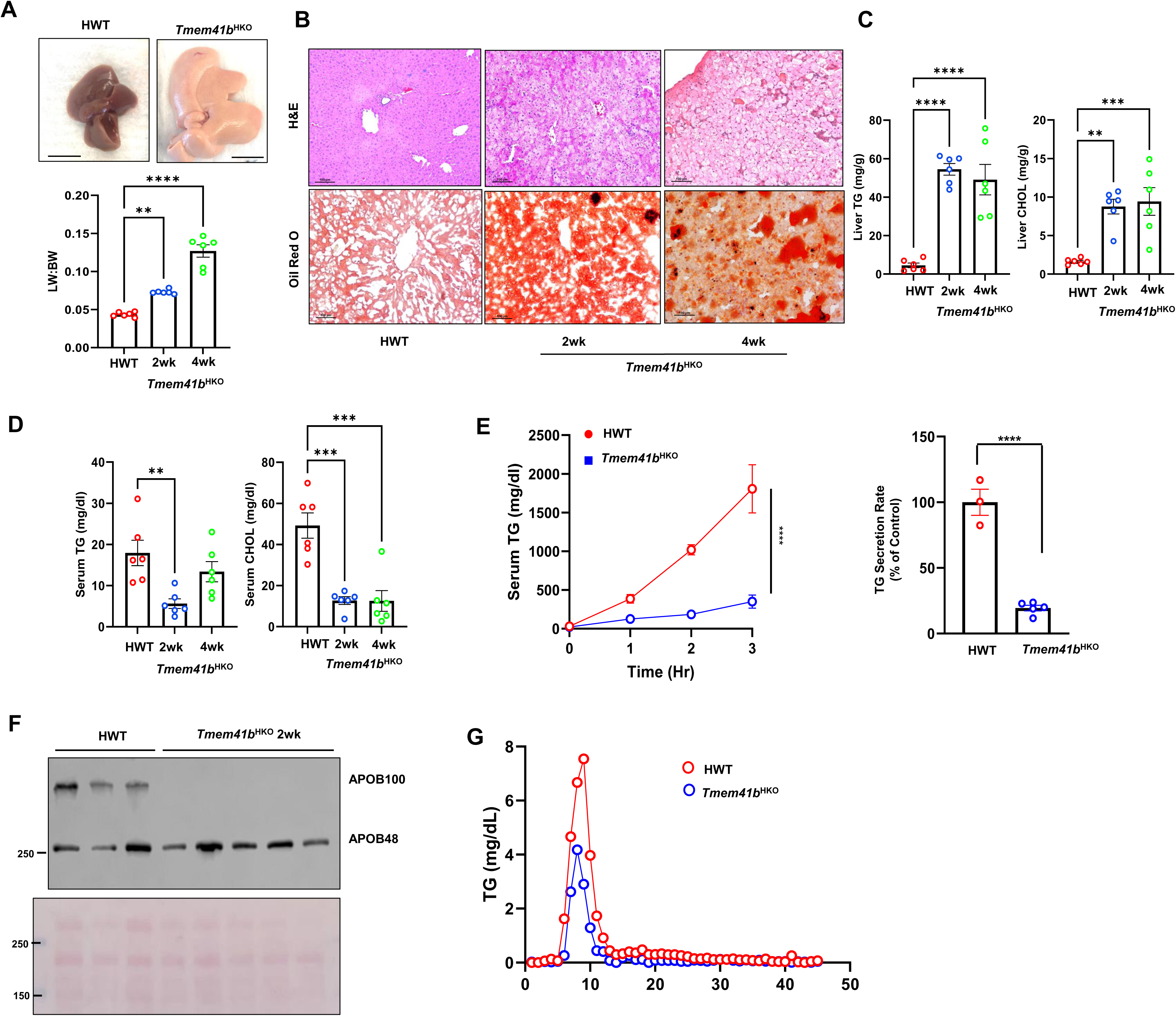
Hepatic deletion of TMEM41B impairs lipoprotein secretion and leads to rapid development of a micro-vesicular steatosis in mice. (A) Representative images of 8-12-week-old *Tmem41b*^flox^ mice at 4 weeks post AAV8-TBG-null (*Tmem41b*^HWT^) or AAV8-TBG-cre (*Tmem41b^HKO^*) injection with liver/body weight ratio. (B) H&E staining of liver tissues from *Tmem41b*^HWT^ and *Tmem41b^HKO^* mice. Scale bars, 100 µm. Hepatic (C) and serum (D) TG and cholesterol (CHOL) were quantified. (E) *Tmem41b*^flox^ mice at 2 weeks post AAV injection were injected with Pluronic™ F-127 and serum TG concentrations were measured. TG secretion rate was quantified. (F) Serum APOB from *Tmem41b*^flox^ mice at 2 weeks post AAV injection were subjected to immunoblot analysis. (G) Mice were fasted 4 hours followed by Pluronic™ F127 injection for another 3 hours. Serum from the same group of mice was pooled (n=4). Lipoprotein profiles were analyzed by FPLC. *p<0.05, **p<0.01, ***p<0.001, ****p<0.0001 (one-way ANOVA with post-hoc Turkey test (A-D) or unpaired Student’s *t test* (E)).

Like *Tmem41b*^HKO^ mice, one-month-old liver-specific*Tmem41b* KO (*Tmem41b*^LKO^) mice also showed hepatomegaly (**Fig. S1A**), massive steatosis (**Fig. S1B**), markedly increased hepatic TG and cholesterol contents (**Fig. S1C**), and decreased serum TG and cholesterol (**Fig. S1D**). VLDL-TG and APOB100 secretion decreased in *Tmem41b*^LKO^ mice, while APOB48 secretion was again preserved (**Fig. S1E**). TG secretion rate was reduced by 66% in *Tmem41b*^LKO^ mice (**Fig. S1E**). Notably, like male mice, female *Tmem41b*^HKO^ mice developed a similar increase in liver-body weight ratio, hepatic TG and cholesterol, and a decrease in serum TG and cholesterol (**Fig. S2A-E**). These data indicate that TMEM41B plays an important role in regulating hepatic APOB100-VLDL secretion and lipid homeostasis in mice, which is consistent with a previous report using guide RNA-mediated gene editing to inactivate *Tmem41b* in mouse livers [11].

### Hepatic deletion of TMEM41B leads to the development of MASH

Serum levels of ALT increased in *Tmem41b*^HKO^ and *Tmem41b*^LKO^ mice (**Fig. 2A, Fig. S3A-B**). Hepatic mRNA levels of inflammatory genes were markedly increased in *Tmem41b*^HKO^ mice (**Fig. 2B**). F4/80-positive macrophages also increased in *Tmem41b*^HKO^ mice (**Fig. 2C**). *Tmem41b*^HKO^ mice displayed increased liver Sirius Red staining at 10 weeks post AAV injection (**Fig. 2D**). Hepatic mRNA levels of fibrotic genes and protein levels of α-smooth muscle actin were elevated in *Tmem41b*^HKO^ mice as early as 2 weeks post AAV injection compared to matched WT mice (**Fig. 2E-F**). Together, these data indicate that hepatic deletion of *Tmem41b* leads to MASH in mice.

**Fig. 2.**
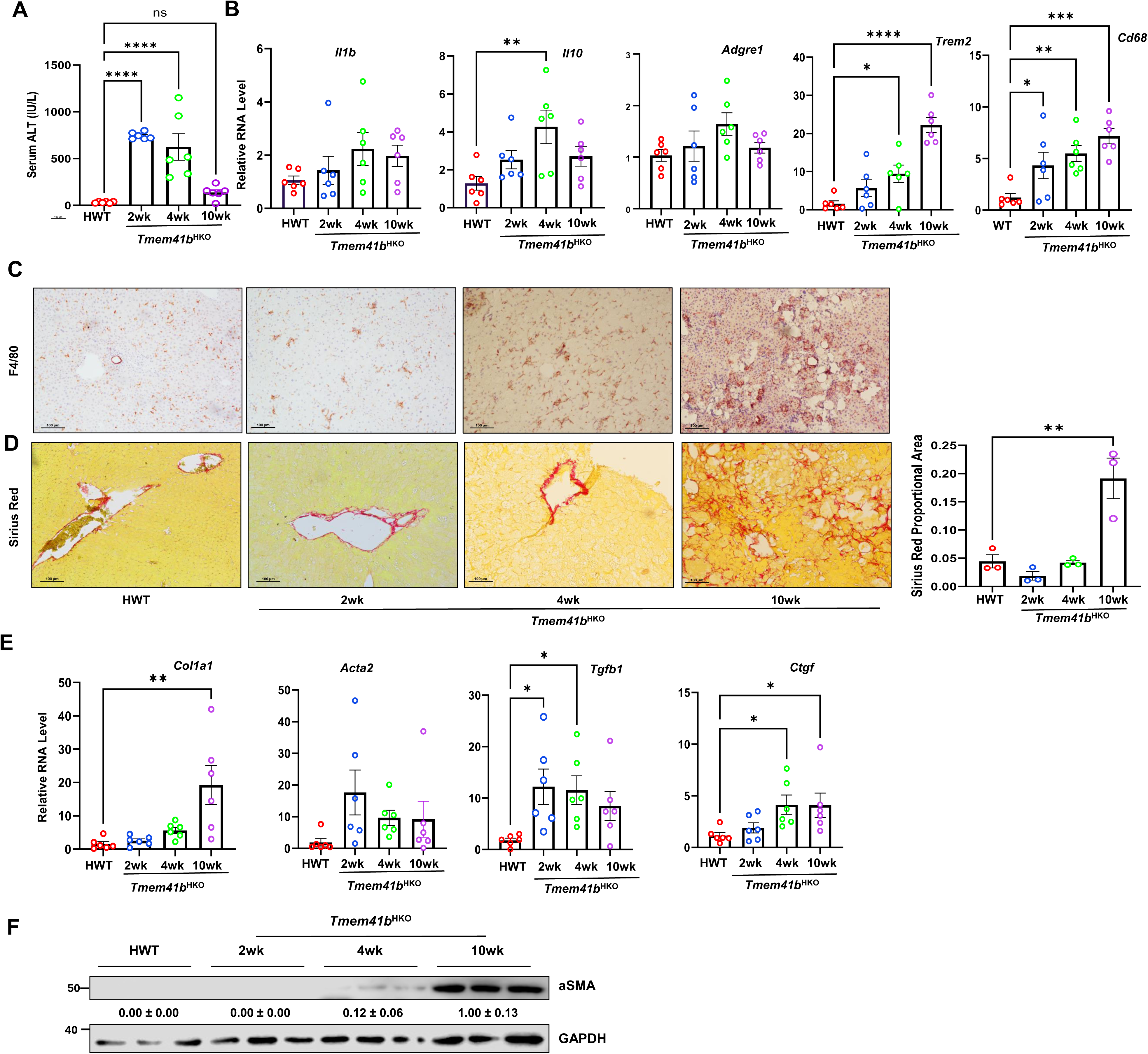
Hepatic deletion of TMEM41B leads to the development of MASH. (A) Serum ALT in *Tmem41b*^flox^ mice at 2, 4 and 10 weeks post AAV injection. (B) Hepatic mRNA levels of pre-inflammatory cytokines and immune cell marker genes were analyzed by qPCR and normalized to *Actb* mRNA. (C) Immunohistochemistry staining for F4/80 in mouse livers. (D) Sirius Red staining in mouse livers. (E) Hepatic mRNA levels of fibrogenic genes were analyzed by qPCR and normalized to *Actb* mRNA. (F) Immunoblot analysis of α-SMA in mouse livers. Data represent mean ± SEM. *p<0.05, **p<0.01, ***p<0.001, ****p<0.0001 (one-way ANOVA with post-hoc Turkey test).

### Decreased TMEM41B is associated with human MASLD and overexpression of TMEM41B alleviates diet-Induced hepatic steatosis

To examine whether TMEM41B is associated with human MASLD, we analyzed the expression of TMEM41B in livers from normal subjects and those with simple steatosis, and MASH. Immunoblot analysis revealed that TMEM41B expression declined in subjects with steatosis and MASH (**Fig. 3A**). H &E staining confirmed increased accumulation of lipid droplets and immune cells in the liver of MASLD (**Fig. 3B**). Mice fed with CDAHFD (choline-deficient, amino acid-defined HFD (45 % fat) containing 0.1% methionine), a MASH diet, also exhibited decreased hepatic TMEM41B (**Fig. 3C**). To examine the therapeutic effect of TMEM41B in CDAHFD-induced steatosis, conditional *Tmem41b* knockin mice were established and fed with CDAHFD (**Fig. 3D**). The overexpression of TMEM41B alleviated CDAHFD-induced steatosis, as shown by improved H&E and Oil Red O staining, an enlarged liver/body weight ratio, and hepatic TG accumulation, but had no effect on elevated serum ALT values. (**Fig. 3E-F**). Overexpression of TMEM41B also restored the decreased TMEM41B caused by CDAHFD in mouse livers without affecting VMP1 expression (**Fig. 3G**). These data indicate that overexpression of TMEM41B improves hepatic steatosis.

**Fig. 3.**
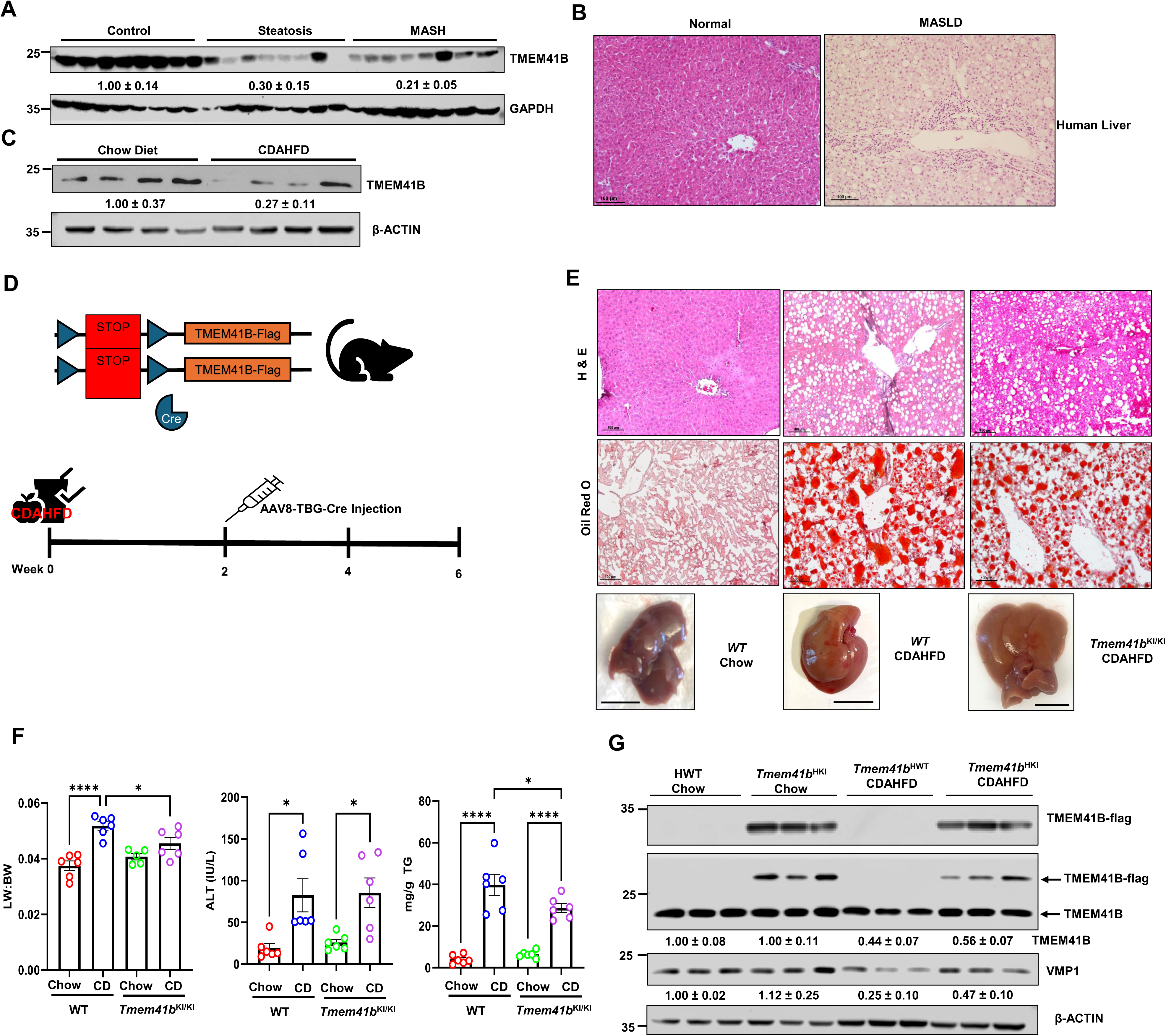
Reduced TMEM41B Is associated with human MASLD livers and overexpression of TMEM41B alleviates diet-Induced hepatic steatosis. (A) Immunoblot analysis of TMEM41B and (B) H&E staining in human livers. (C) Protein level of TMEM41B in mice fed with CDAHFD. (D) Scheme of CDAHFD-induced MASH in mice. (E) Representative H&E and Oil O Red images from mice. Scale bars, 50 µm. (F) Liver-body weight ratio, ALT and hepatic TG in CDAHFD-fed mice. (G) Immunoblot analysis of TMEM41B and VMP1 in mice fed with CDAHFD. Data represent mean ± SEM. * p<0.05; **** p<0.0001 (one-way ANOVA with post-hoc Turkey test).

### Hepatic deletion of *Tmem41b* or *Vmp1* leads to decreased hepatic levels of phospholipids and disrupts ER-mitochondria contact

Since the deletion of VMP1 in the liver causes impaired VLDL secretion and MASH in mice [16], we proceeded to compare the effects of deleting either VMP1 or TMEM41B on changes in hepatic phospholipids, and the ultrastructure of the ER-mitochondria contact, as well as the ER and Golgi [16]. Unbiased lipidomic analysis revealed that the levels of phosphatidylethanolamine (PE), but not phosphatidylcholine (PC), decreased significantly in *Tmem41b* ^HKO^ mice livers. However, both PC and PE decreased significantly in *Vmp1*^HKO^ mouse livers. Neutral lipids, including cholesteryl esters (CE), diacyl- and triacylglycerols increased dramatically in *Tmem41b*^HKO^ or *Vmp1*^HKO^ mouse livers (**Fig. 4A**). However, the magnitude of these changes appeared relatively more pronounced in mice lacking hepatic *Vmp1* compared to those lacking *Tmem41b*. There were minimal changes in the hepatic metabolites of fatty acid synthesis, medium chain fatty acid and long chain saturated and polyunsaturated fatty acid as well as various acyl carnitine fatty acid in *Tmem41b*^HKO^ mice, but these metabolites were markedly increased in *Vmp1*^HKO^ mice (**Fig S4 and S5**). The acyl chain length and levels of saturation of phospholipids play critical roles in membrane trafficking processes [24]. In the livers of *Tmem41b^HKO^*mice, the levels of shorter acyl chain PC species decreased following AAV-TBG-Cre injection after 2 weeks, but not after 1 week. In contrast, the decrease of shorter acyl chain PC species in *Vmp1^HKO^* mouse livers was more significant and occurred as early as 1 week post-injection. However, longer acyl chain PC species exhibited a more substantial increase in *Tmem41b^HKO^* mouse livers compared to *Vmp1^HKO^*mouse livers (**Fig. S6**). For PE species, both shorter and longer acyl chain species decreased in T *Tmem41b^HKO^* mouse livers post-AAV-TBG-Cre injection after 2 weeks but not after 1 week. In *Vmp1^HKO^*mouse livers, both shorter and longer-chain PE species decreased after AAV-TBG-Cre injection at both 1 week and 2 weeks (**Fig. S7**). Overall, the reductions in both PC and PE species were more pronounced in *Vmp1^HKO^* HKO mouse livers than in *Tmem41b^HKO^* mouse livers.

**Fig. 4.**
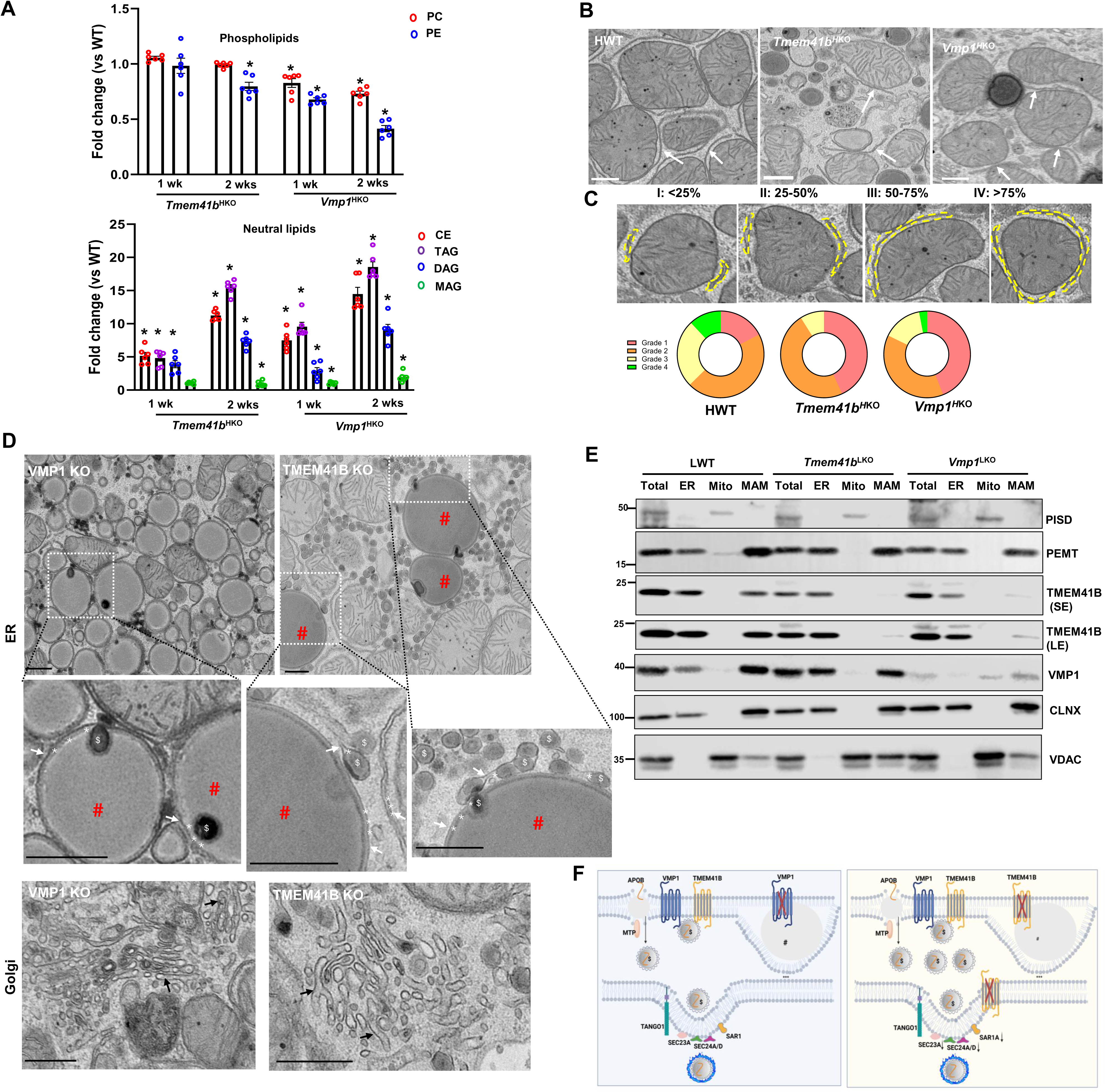
Hepatic deletion of *Tmem41b* or *Vmp1* leads to a decline in phospholipids and disrupts ER-mitochondria contact. (A). Total phospholipids and neutral lipids of mouse livers from *Tmem41b*^HKO^ and Vmp1^HKO^ mice by lipidomic analysis (n=6). (B) Representative EM images from *Tmem41b*^HKO^ and Vmp1^HKO^ mice (2 weeks post AAV8 injection). White Arrows denote close contact of ER with mitochondria approximately< 30 nm, which are defined as MAM. (C) ER-mitochondria contact sites were quantified in mouse livers from at least 10 images. Scale bars, 500 nm. (D) Representative EM images of lipids from *Tmem41b*^HKO^ and Vmp1^HKO^ mice (2 weeks post AAV8 injection). White arrows denote phospholipid bilayers of the ER. Stars denote ER lumen, # represents LDs, $ represents lipoproteins. (E) Subcellular fractions from 1-month-old LWT, *Tmem41b*^LKO^ and *Vmp1*^LKO^ mice were subjected to immunoblot analysis with the indicated antibodies. (F) Diagrams of lipid droplets in hepatic VMP1 and TMEM41B KO mouse hepatocytes. Data represent mean ± SEM. *p<0.05 (unpaired Student’s *t test*).

ER-Mitochondria contact sites are critical for phospholipid synthesis and calcium transport [25]. EM analysis revealed that ER-Mitochondria contact sites were readily detected in HWT, *Tmem41b*^LKO^, and *Vmp1*^LKO^ mouse hepatocytes (**Fig. 4B**). ER-Mitochondria contact sites were graded based on a 4-point scale to assess the contact between ER and mitochondria, with Grade 1 being minimal contact (<25%) and Grade 4 indicating mitochondria nearly completely enveloped by ER (>75%). Quantitative EM analysis showed fewer ER-Mitochondria contact sites in *Tmem41b*^LKO^ or *Vmp1*^LKO^ mouse hepatocytes than WT hepatocytes (**Fig. 4C**). The loss of these contact sites seemed to be more pronounced in TMEM41B-deficient hepatocytes than in VMP1-deficient hepatocytes.

Consistent with our previous report [16], the “LDs” observed in VMP1 KO hepatocytes exhibited membranes that are likely phospholipid bilayers of the ER, facing the cytosol (indicated by white arrows). These LDs displayed clear electron-dense edges, which probably represent the surrounding phospholipid monolayer around the lipid structures. The spaces marked by stars between the ER membrane and the electron-dense-edged lipid structures represent the ER lumen, suggesting that these LDs are stalled at the ER phospholipid bilayer (**Fig 4D, “#”**). Interestingly, a small portion of the LDs was found within the ER lumen, which is likely composed of lipoproteins (denoted as “$”). Similar phospholipid bilayer LDs were also identified in TMEM41B KO hepatocytes (**Fig 4D, “#”).** However, the number of ER luminal LDs (lipoproteins) was considerably higher in TMEM41B KO hepatocytes compared to VMP1 KO hepatocytes. Since lipoproteins were either stalled at the ER phospholipid bilayer or located inside the ER lumen, the Golgi apparatus in both VMP1 and TMEM41B KO hepatocytes was entirely devoid of lipoproteins (**Fig 4D, black arrows**).

To further investigate the possible effects of *Tmem41b* or *Vmp1* KO on ER-mitochondria contact, cell fractionation was performed using livers from hepatic *Tmem41b* or *Vmp1* KO and their WT littermates. Both TMEM41B and VMP1 were enriched at the ER and mitochondrial-associated membrane (MAM) (**Fig. 4E**). Phosphatidylethanolamine N-methyltransferase (PEMT) and phosphatidylserine decarboxylase (PISD) are two enzymes important for PC and PE synthesis. While PEMT was enriched at the ER and MAM with higher levels on MAM in WT mouse livers, *Tmem41b*^LKO^ or *Vmp1*^LKO^ mice had reduced total PEMT and PEMT in MAM fractions (**Fig. 4E**). PISD was only located on mitochondria with no change or slight decrease in hepatic level in *Tmem41b*^LKO^ or *Vmp1*^LKO^ mice (**Fig. 4E**). VMP1 protein level was reduced at MAM in *Tmem41b*^LKO^ mouse livers. Conversely, loss of VMP1 led to a profound decrease in TMEM41B localization to MAM sites. However, considerable levels of TMEM41B were detected in the ER fraction of *Tmem41b*^LKO^ mouse livers, which could be from non-parenchymal cells. To our surprise, a trace of VMP1 was observed on mitochondria fractions and need further investigation (**Fig. 4E**). Together, both VMP1 and TMEM41B localize on the MAM, and the loss of either results in decreased hepatic phospholipid content and stalled lipid secretion at ER phospholipid bilayer and the ER lumen (**Fig. 4F**). Loss of VMP1 appears to have more profound effects on the changes of hepatic phospholipids and fatty acid metabolism than loss of TMEM41B.

### Knockout of both *Tmem41b* and *Vmp1* does not further exacerbate impaired lipoprotein secretion and hepatic steatosis

To further assess the relationship between TMEM41B and VMP1 in regulating VLDL secretion, hepatic *Tmem41b* /*Vmp1* double KO (DKO) mice were generated. DKO mice developed steatosis and hepatomegaly, similar to *Tmem41b*^LKO^ or *Vmp1*^LKO^ mice (**Fig. 5A-C**). *Vmp1*^LKO^ and DKO mice had similar serum ALT levels that were higher than those of *Tmem41b*^LKO^ mice (**Fig. 5C**). Hepatic TG and cholesterol increased to a similar extent in all mice (**Fig. 5C**). *Vmp1*^LKO^ and DKO mice had more severe VLDL secretion defects compared to *Tmem41b*^LKO^ mice. TG secretion rate was reduced by 49.44%, 62.88%, and 69.38%, respectively, in *Tmem41b*^LKO^ and *Vmp1*^LKO^, and DKO mice (**Fig. 5D**). Expression of MTTP and disulfide isomerase (PDI), both of which are required for VLDL assembly, were increased in *Tmem41b*^LKO^ and DKO but not in *Vmp1*^LKO^ mice (**Fig. 5E**). While several COPII proteins, SEC23A, SEC24A, and SEC24D, required for exit of VLDL from ER, were moderately reduced in *Vmp1*^LKO^ mouse livers, their protein levels were decreased profoundly in either *Tmem41b*^LKO^ or *DKO* mouse livers (**Fig. 5E**). Interestingly, while loss of TMEM41B did not change VMP1 protein levels, loss of VMP1 profoundly decreased TMEM41B protein in mouse livers (**Fig. 5E**). PEMT and PISD were profoundly decreased in *Vmp1*^LKO^ mouse livers, which was not further decreased in DKO livers (**Fig. 5F**). These results suggest that TMEM41B and VMP1 have different effects on the proteins that regulate VLDL secretion. In general, the loss of VMP1 appears to have a more significant impact, leading to greater impairment of VLDL secretion and increased liver injury.

**Fig. 5.**
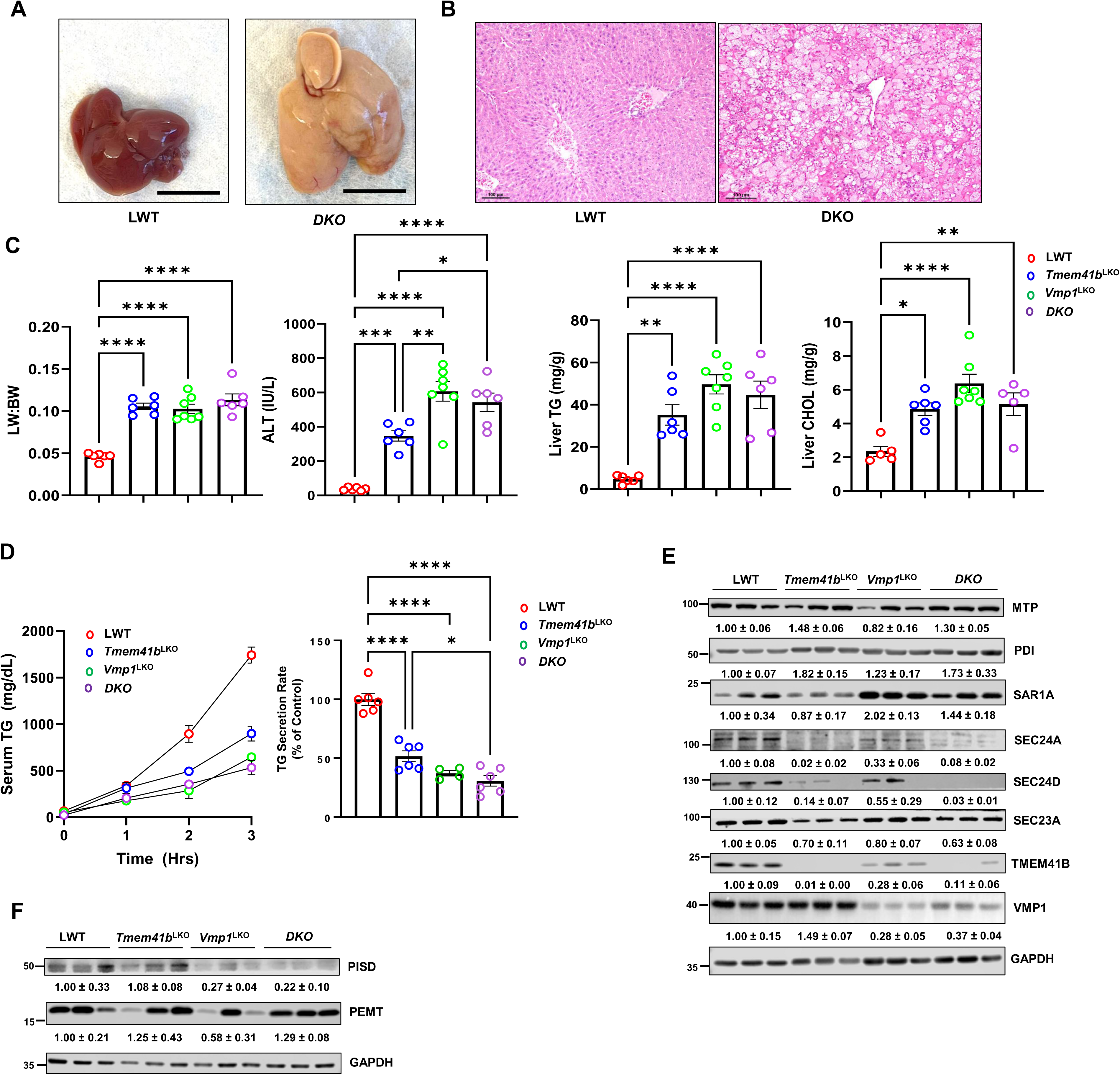
Knockout of both *Tmem41b* and *Vmp1* does not further exacerbate impaired lipoprotein secretion and hepatic Steatosis. (A) Representative images of 1-month-old WT and DKO mice. Scale bars, 1cm. (B) H&E staining of liver tissues from the mice. Scale bars, 100 µm. (C) Liver/body weight ratio, ALT, and hepatic TG and cholesterol (CHOL) were measured in mice. (D) LWT, *Tmem41b^LKO^*, *Vmp1^LKO^*, and DKO mice were injected with Pluronic™ F-127 and serum TG concentrations were measured. TG secretion rate was quantified. (E) Total liver lysates were extracted from the mice. Total mouse liver lysates were subjected to immunoblot analysis for the indicated proteins and followed by densitometric analysis. (F) PEMT and PISD were examined by immunoblots. Data represent mean ± SEM. *p<0.05, ***p<0.001, ****p<0.0001 (one-way ANOVA with post-hoc Turkey test).

### Overexpression of hepatic VMP1 or TMEM41B partially alleviated steatosis caused by the loss of the other in mice

To explore the relationship between VMP1 and TMEM41B in VLDL secretion, we examined whether overexpression of either VMP1 or TMEM41B could compensate for the deficiency of the other on hepatic lipid homeostasis. Overexpression of VMP1 significantly improved pale liver color in *Tmem41b*^LKO^ mice and reduced microsteatosis, as observed through H&E staining of liver sections. There was no notable difference in steatosis improvement between heterozygous and homozygous overexpression of VMP1 (**Fig. 6A-B**). Additionally, both heterozygous and homozygous overexpression of VMP1 partially corrected elevated levels of serum ALT and liver-to-body weight ratio in *Tmem41b*^LKO^ mice (**Fig. 6C**). Overexpression of VMP1 also resulted in a significant reduction of hepatic TG and cholesterol in *Tmem41b*^LKO^ mice, regardless of whether VMP1 was overexpressed as a heterozygous or homozygous allele (**Fig. 6D**).

**Fig. 6.**
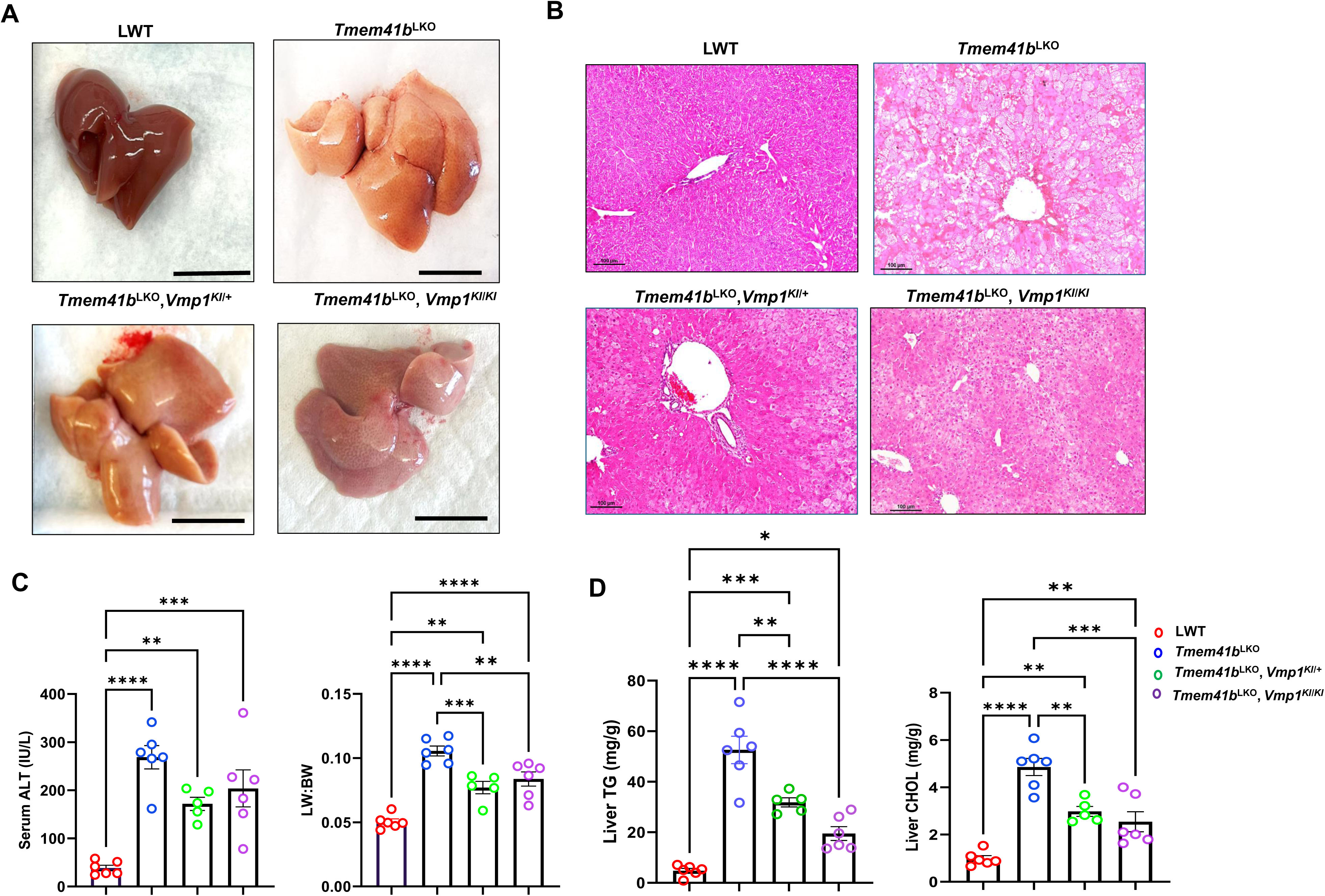
Overexpression of hepatic VMP1 partially alleviated steatosis caused by the loss of TMEM41B. (A) Representative images of 1-month-old *Tmem41b*^LKO^, *Tmem41b*^LKO^/*Vmp1^KI^*^/+^ and *Tmem41b*^LKO^/*Vmp1*^KI/KI^ mice. Scale bars, 1cm. (B) H&E and Oil O Red staining of liver tissues from the mice. Scale bars, 100 µm. (C) Liver/body weight ratio and ALT were measured. (D) Hepatic TG and cholesterol (CHOL) were measured in mice. Data represent mean ± SEM. *p<0.05, **p<0.01, ***p<0.001, ****p<0.0001 (one-way ANOVA with post-hoc Turkey test).

Overexpression of TMEM41B in *Vmp1*^LKO^ mice alleviated steatosis as demonstrated by H&E staining of liver sections (**Fig. 7A-B**). Overexpression of TMEM41B also partially corrected increased liver body weight ratio and serum levels of ALT in *Vmp1*^LKO^ mice (**Fig. 7C**). Levels of hepatic TG and cholesterol were also reversed with overexpression of TMEM41B in *Vmp1*^LKO^ mice (**Fig. 7D**). Notably, homozygous *Tmem41b* KI mice had much better improvement than heterozygous *Tmem41b* KI mice for the above parameters in *Vmp1*^LKO^ mice. The restoration of TMEM41B in *Vmp1*^LKO^ mice also improved VLDL secretion. TG secretion rate was reduced by 69.87% in *Vmp1*^LKO^ mice but improved by 21.7% with TMEM41B overexpression (**Fig. 7E**). Together, these data indicate that VMP1 and TMEM41B functionally interact and partially compensate for the loss of the other on hepatic VLDL secretion.

**Fig. 7.**
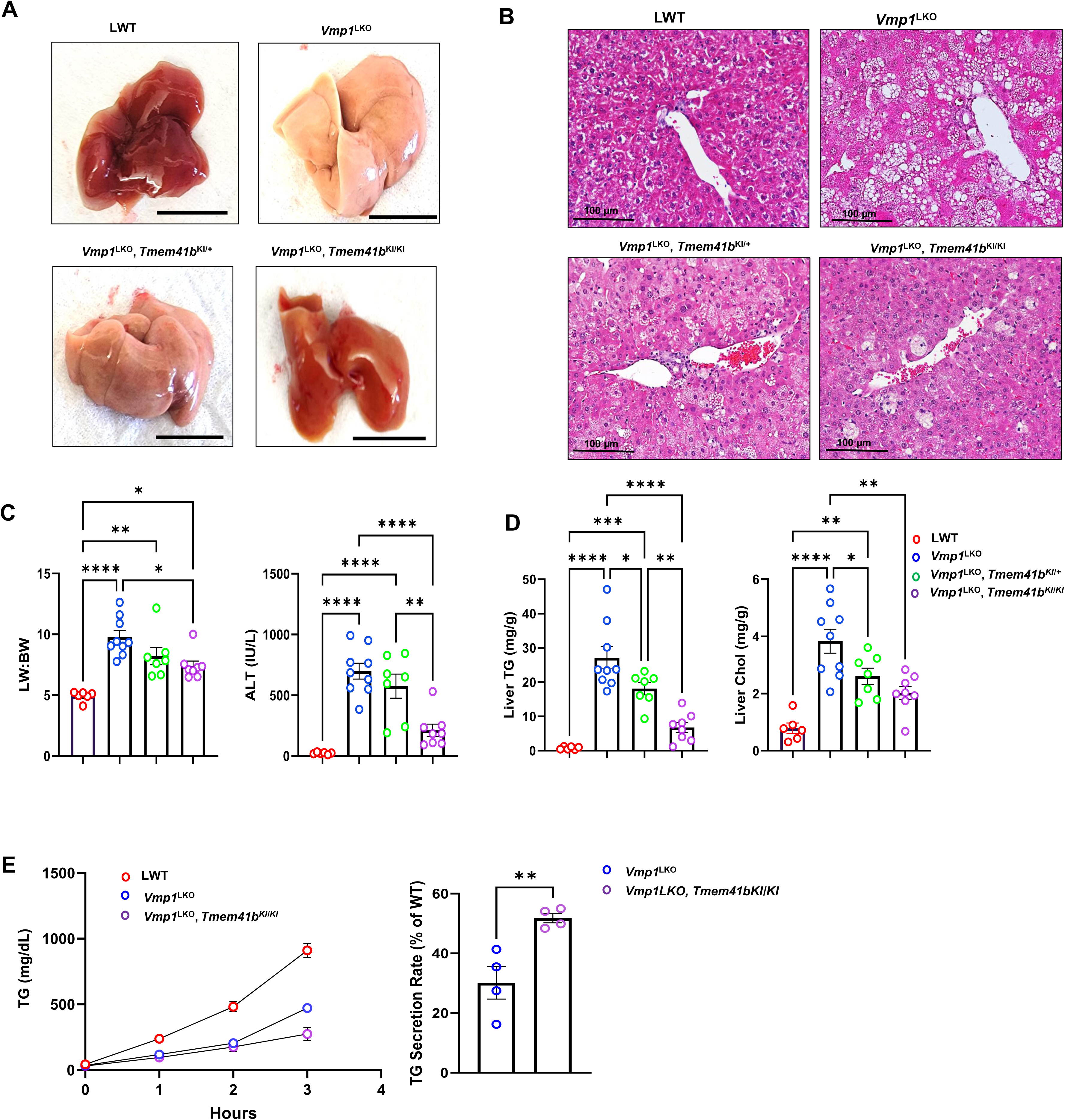
Overexpression of hepatic TMEM41B improved VLDL secretion and partially alleviated Steatosis caused by the loss of VMP1. (A) Representative images of 1-month-old *Vmp1*^LKO^, *Vmp1*^LKO^/*Tmem41b*^KI/+^ *and Vmp1*^LKO^/*Tmem41b*^KI/KI^ mice. Scale bars, 1cm. (B) H&E staining of liver tissues from the mice. Scale bars, 100 µm. (C) Liver/body weight ratio and ALT were measured. (D) Hepatic TG and cholesterol (CHOL) were measured in mice. (E) LWT, *Vmp1*^LKO^, *Vmp1*^LKO^/*Tmem41b*^KI/+^ *and Vmp1*^LKO^/*Tmem41b*^KI/KI^ mice were injected with Pluronic™ F-127 and serum TG concentrations were measured. VLDL secretion rates were quantified. Data represent mean ± SEM. *p<0.05, **p<0.01, ***p<0.001, ****p<0.0001 (one-way ANOVA with post-hoc Turkey test).

### Overexpression of TMEM41B in *Vmp1*^LKO^ and VMP1 in *Tmem41b*^LKO^ mice partially corrects autophagy defect

Consistent with previous findings [26], immunoblot analysis showed that levels of LC3B-II and p62 were increased in *Tmem41b*^HKO^ mouse livers (**Fig. 8A**), indicating impaired autophagy by deletion of *Tmem41b* in mouse livers. The levels of LC3-II and p62 were comparable in *Vmp1*^LKO^ *and DKO* mice but were much higher than those of *Tmem41b*^LKO^ mice (**Fig. 8B**), suggesting VMP1 may be more critical than TMEM41B in regulating autophagy in mouse livers. Consistent with this, the levels of hepatic dipeptides generated by autophagic lysosomal degradation decreased in *Tmem41b*^LKO^ and *Vmp1*^LKO^ mice, with a more pronounced decrease observed in *Vmp1*^LKO^ mice compared to *Tmem41b*^LKO^ mice (**Fig. S8**). We next examined whether overexpression of either VMP1 or TMEM41B could compensate for the deficiency of the other on hepatic autophagy in mouse livers. Heterozygous overexpression of VMP1 completely corrected increased levels of hepatic LC3-II but not p62 in *Tmem41b*^LKO^ mouse livers (**Fig. 8C**). Homozygous overexpression of VMP1 further increased LC3B-II and p62 compared with heterozygous overexpression of VMP1 **(Fig. 8C)**. Levels of LC3B-II and p62 were markedly increased in *Vmp1*^LKO^ mouse livers compared with matched WT mouse livers. Heterozygous overexpression of TMEM41B partially corrected increased levels of hepatic LC3-II and p62 in *Vmp1*^LKO^ mouse livers, which were further corrected by homozygous overexpression of TMEM41B (**Fig. 8D**). These results suggest that loss of VMP1 has more profound defects on hepatic autophagy than loss of TMEM41B. TMEM41B and VMP1 may partially compensate for the loss of the other in hepatic autophagy; however, a precise level of VMP1 is critical for effective autophagy in hepatocytes.

**Fig. 8.**
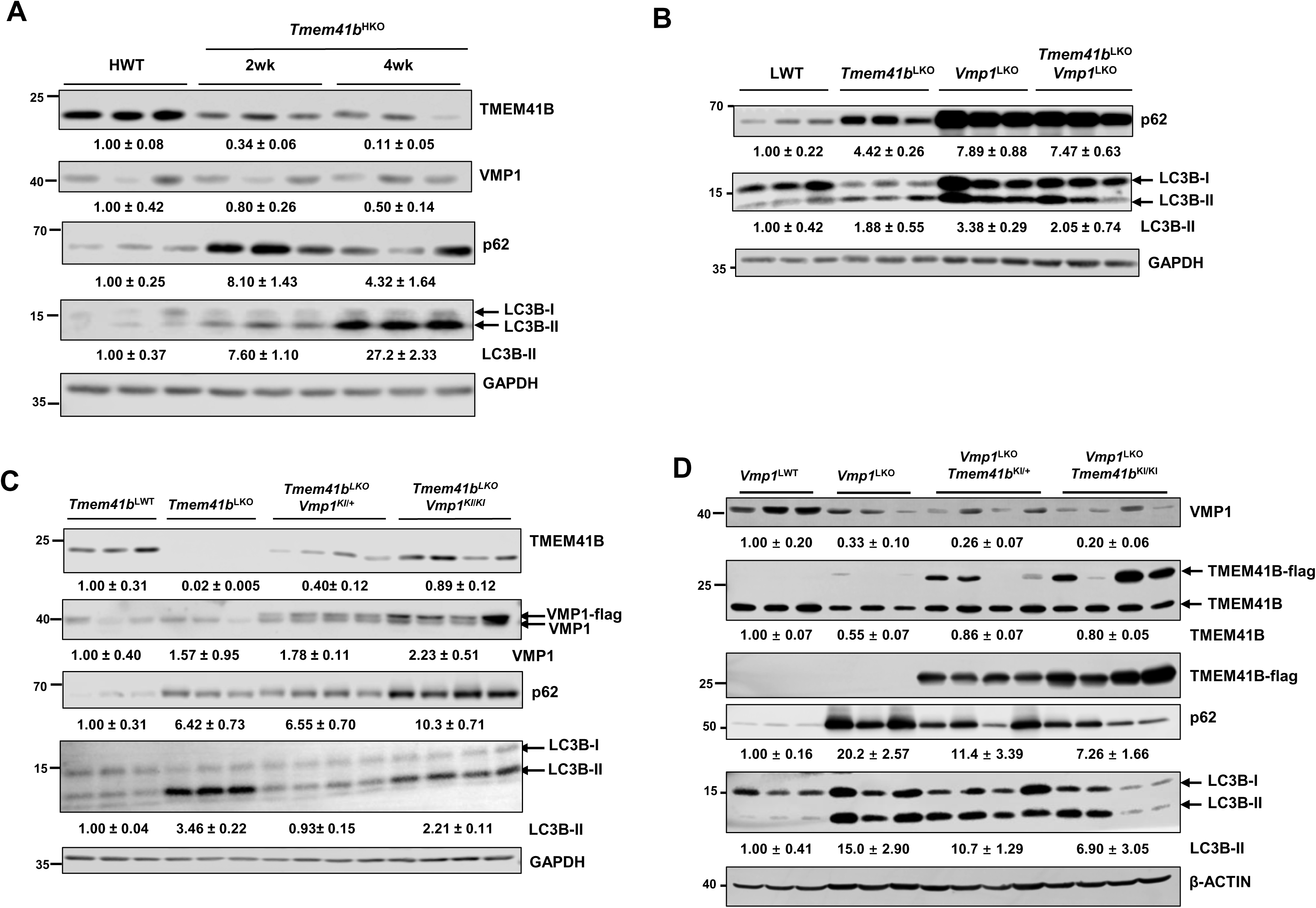
Overexpression of TMEM41B in *Vmp1*^LKO^ and VMP1 in *Tmem41b*^LKO^ mice partially corrects autophagy defect. (A) Total lysates were extracted from *Tmem41b*^flox^ mice at 2 and 4 weeks post AAV injection, and subjected to the immunoblot analyses with the indicated antibodies followed by densitometric analysis. (B) Total lysates were extracted from LWT, *Tmem41b*^LKO^, *Vmp1*^LKO^, and DKO mice and subjected to immunoblot analyses with the indicated antibodies followed by densitometric analysis. (C) Total lysates were extracted from LWT, *Tmem41b*^LKO^, *Tmem41b*^LKO^/*Vmp1*^KI/+^ and *Tmem41b*^LKO^/*Vmp1*^KI/KI^ mice and subjected to the immunoblot analyses with the indicated antibodies followed by densitometric analysis. (D) Total lysates were extracted from LWT, *Vmp1*^LKO^, *Vmp1*^LKO^/*Tmem41b*^KI/+^ *and Vmp1*^LKO^/*Tmem41b*^KI/KI^ mice and subjected to the immunoblot analyses with the indicated antibody followed by densitometric analysis.

## Discussion

In this study, we found that *Vmp1*^LKO^ mice developed more severe VLDL secretion defects and liver injury than *Tmem41b*^LKO^ mice, and that hepatic deletion of both *Vmp1* and *Tmem41b* did not further impair VLDL secretion and worsen liver injury compared with *Vmp1*^LKO^ mice. *Vmp1*-deficient hepatocytes exhibited increased ER bilayer “LD” and fewer ER luminal “LD,” while *Tmem41b*-deficient hepatocytes showed increased ER bilayer “LD” and a greater number of ER luminal “LD.” Overexpression of *Vmp1* or *Tmem41b* in *Tmem41b*-deficient or *Vmp1*-deficient mouse livers partially alleviated steatosis and corrected autophagy defects, suggesting TMEM41B and VMP1 functionally interact and compensate for hepatic VLDL secretion and autophagy. However, the restoration of VLDL secretion and hepatic autophagy by *Tmem41b* in in *Vmp1*-deficient mouse livers is in a gene dose-dependent manner. In contrast, overexpression of either low or high levels of *Vmp1* in *Tmem41b*-deficient mouse livers partially improved VLDL secretion and hepatic steatosis, but high levels of Vmp1 fail to rescue hepatic autophagy defects, implying that VMP1 and TMEM41B have similar yet distinct roles in regulating hepatic VLDL secretion and autophagy. Previous studies that compared predicted TMEM41B and VMP1 structures have identified sequence homology between these two proteins. Notably, both VMP1 and TMEM41B contain a DedA domain consisting of two transmembrane helices, one extramembrane helix, and two reentrant loops. TMEM41B and VMP1 seem to structurally differ primarily in their N-termini, in which TMEM41B contains fewer transmembrane domains, but an extramembrane domain is not present in VMP1 [27, 28]. Whether these structural differences would contribute to the differences in their regulation in hepatic VLDL secretion, liver injury, and autophagy remains to be investigated in the future.

Conventional LDs are cytosolic organelles that store metabolic energy and consist of a hydrophobic core of neutral lipids surrounded by a phospholipid monolayer decorated with proteins. Histological analysis with H&E staining of liver sections revealed a significant increase in the accumulation of hepatic LDs in both *Tmem41b*-deficient and *Vmp1*-deficient mouse livers. EM analysis of the ultrastructure of these LDs in VMP1 KO hepatocytes showed that the large LDs appear to be stalled within the ER membrane bilayer, consistent with our previous findings [16]. Additionally, we observed some small LDs inside the ER lumen, surrounded by ER membranes. Interestingly, we also identified hybrid LDs, where large ER membrane bilayer LDs coexist with smaller luminal LDs in the same ER structure. These ER luminal LDs are more prominent in TMEM41B KO hepatocytes compared to VMP1 KO hepatocytes. These small ER luminal LDs likely represent lipoproteins transported from the ER bilayer into the ER lumen for further lipidation and maturation. A previous study, which utilized a gRNA against *Tmem41b* and AAV-TBG-Cre injection in mice harboring the lox-STOP-lox Cas9 cassette, demonstrated that TMEM41B-deficient hepatocytes lack mature lipoproteins with increased LDs encapsulated by highly curved ER membranes [11]. These data suggest that TMEM41B and VMP1 are necessary for VLDL biogenesis in the ER.

How do TMEM41B and VLDL regulate VLDL biogenesis, and what differences exist between them? Three processes critical for VLDL secretion occur at specific ER sites. These include (1) import of neutral lipids from the ER bilayer into the ER lumen, (2) assembly of pre-VLDL and maturation in the ER lumen, and (3) export of pre-VLDL from the ER lumen. ER membrane phospholipid content and composition are critical for ER membrane remodeling, curvature and tension, which are also crucial for LD budding from the ER bilayer [29, 30]. The amount of PC and PE as well as the fatty acyl chain compositions of PC and PE, especially PC and PE arachidonyl chain, has been shown to regulate VLDL secretion [31, 32]. The decreased hepatic PC and PE content, together with the changes to the acyl-chain composition of PC and PE in TMEM41B or VMP1 KO hepatocytes may thus change the biophysical tension and curvature of the ER membrane, arresting import of neutral lipids from the ER membrane bilayer to the ER lumen. VMP1 KO hepatocytes have a more profound decrease of total PL and short- and long-chain PC and PE species than TMEM41B KO hepatocytes, which may explain the greater number of LDs stalled in the ER membrane bilayer than TMEM41B KO hepatocytes. MAM is critical in regulating PL synthesis and transfer between the ER and mitochondria. TMEM41B KO and VMP1 KO hepatocytes have decreased ER-mitochondria contact sites. Furthermore, both TMEM41B and VMP1 are located at MAM, and PEMT and PISD, two necessary PL synthesis enzymes, were reduced in TMEM41B and VMP1 KO livers, which may contribute to reduced PL synthesis. While TMEM41B and VMP1 are both ER scramblases that regulate the PL distribution between the outer and inner leaflet of the ER membrane [12, 27, 33], it remains unclear whether TMEM41B and VMP1 would have different scramblase activity. Both TMEM41B and VMP1 possess a DedA domain and were structurally predicted to serve as half transporters for ion and lipid transport [34]. Future work to dissect the structural differences of these two similar proteins may yield further mechanistic insights to better understand how they regulate the ER membrane dynamics and VLDL biogenesis.

Inside the ER lumen, it is known that MTTP is required to transfer the bulk of triglycerides into APOB100 for VLDL assembly. Deletion of MTTP abolishes VLDL secretion. MTTP activity is enhanced by PDI, which is regulated by the IRE1α-XBP1s-PDI axis, and decreased IRE1α-mediated PDI production impairs hepatic VLDL secretion in mice [35–37]. Pre-VLDL particles are progressively lipidated, which requires MTTP and other players, including TM6SF2 [38]. Liver-specific *Tm6sf2* KO mice exhibit steatosis and reduced VLDL secretion with small, underlipidated VLDL particles without affecting APOB48 secretion. We did not find significant changes in either hepatic MTTP or PDI in *Tmem41b* or *Vmp1* KO hepatocytes, suggesting that alterations in MTTP and PDI are less likely to contribute to steatosis in these KO mice. Assembly of pre-VLDL in the ER lumen is APOB-dependent [39]. It has been reported that TMEM41B and VMP1 directly interact with APOB100 [11, 40]. We found that hepatic APOB100 levels markedly decreased in hepatic *Tmem41b* or *Vmp1* KO mouse serum, which may indicate preferential impacts on pre-VLDL assembly in the ER lumen impairing VLDL secretion [41].

At the ER exit site, it is known that TANGO1 and TALI are critical in regulating the export of the pre-VLDL from the ER. Moreover, the COPII-coated vesicle is assembled through a signaling pathway beginning with SAR1 and SURF4 and progressing through SEC proteins at the ER exit site to facilitate delivering pre-VLDL to the Golgi for further processing. It has been reported that TMEM41B interacts with SURF4 at ER exit sites to assist in COP-II-coated vesicle formation [11]. VMP1 also interacted with SEC24D and substantially colocalized with SEC24D in hepatocytes of mouse livers. TMEM41B and VMP1 may likely increase the stability of APOB and COPII complex proteins [14]. However, more profound decreases of SAR1A, SEC24A, SEC24D and SEC23A were found in TMEM41B KO mouse livers than VMP1 KO mouse livers, suggesting that TMEM41B seems to be a more important regulator of COPII-coated vesicle transport at the ER exit sites than VMP1. Future studies are needed to dissect how TMEM41B and VMP1 might regulate hepatic APOB and COPII proteins at either posttranslational, transcriptional, or both. Notably, it has been reported that VMP1 and TMEM41B interact with each other. [26]. As deletion of VMP1 decreased TMEM41B levels, TMEM41B and VMP1 potentially stabilize each other within hepatocytes.

In addition to their function in VLDL secretion, TMEM41B and VMP1 also regulate autophagy at an early step likely by promoting autophagosome closure [26, 42, 43]. Like in a previous report [26], we found autophagy defect to be more severe in VMP1^LKO^ mouse livers than in TMEM41B^LKO^ livers. Hepatic deletion of both TMEM41B and VMP1 did not further exacerbate impaired autophagy. We also found that low-level VMP1 overexpression partially corrected autophagy defect in VMP1-deficient mouse livers. However, high-level VMP1 overexpression impaired autophagy in TMEM41B-deficient mouse livers, suggesting that too much VMP1 may also impair autophagy, though the mechanisms behind this hormesis dose response need further study. TMEM41B could correct autophagy defects in VMP1^LKO^ mouse livers in a dose-dependent manner, which differs from findings from a previous report using cultured HEK293 cells [26]. Likely, the fine-tuned TMEM41B and VMP1 levels in mouse hepatocytes *in vivo* could be different from those of cultured HEK293 cells. Notably, several *in vitro* studies showed that VMP1 deficiency causes an increase in membrane contact sites such as ER-mitochondria, ER-endosomes, ER-lipid droplets, and ER-autophagosomes through VAP-mediated or other mechanisms, suggesting that VMP1 inhibits contact site formation in HeLa, COS7, and MEF cells [13, 43, 44]. In contrast to VMP1, the involvement of TMEM41B in membrane contact has not been well investigated. We and others found that ER-mitochondria contact sites or perhaps total ER were reduced in *Tmem41b*- or *Vmp1*-deficient hepatocytes [11, 14]. It seems organelle contact regulation by VMP1 and TMEM41B may be cell-type dependent. However, how TMEM41B and VMP1 regulate ER-mitochondria and other organelle contact sites, along with their roles in regulating autophagy, needs further investigation.

A recent report identified rare and low-frequency ATG7 loss-of-function variants that promote MASLD progression by impairing autophagy and facilitating ballooning and inflammation in the European population [45]. Increased cell death, inflammation, and fibrosis have been observed in hepatic Atg5 and Atg7-deficiency mice, which have impaired autophagy, but not impaired VLDL secretion and steatosis [34]. However, most genetically modified mouse MASLD models, such as hepatic *Surf4-, Lpcat3-*, *Mea6-*, and *Sar1b*-deficiency mice, showed defective hepatic VLDL secretion without altered AOPB100 and hepatic steatosis, though none of them developed MASH [31, 46–48]. Therefore, the MASH phenotypes in VMP1 and TMEM41B KO mice are distinct from those of MASLD models. The MASH phenotypes in *Vmp1* and *Tmem41b* KO mice are likely due to combined impaired VLDL secretion and autophagy, which is unique to TMEM41B and VMP1.

In summary, we found that human MASH livers have decreased hepatic VMP1 and TMEM41B. Our results indicate that the lack of hepatic TMEM41B and VMP1 impairs VLDL secretion and autophagy, resulting in MASH in mice. Overexpression of VMP1 alleviates steatosis but not autophagy at high expression levels, whereas overexpression of TMEM41B improves VLDL secretion, ameliorates steatosis, and partially corrects autophagy defects. TMEM41B and VMP1 likely have both redundant and distinct roles in regulating VLDL secretion and autophagy, and future studies designed to boost hepatic TMEM41B but not VMP1 may be beneficial in enhancing VLDL secretion and improving MASH pathology in humans.

## ACKNOWLEDGEMENTS

The authors acknowledge KUMC Liver Center for human liver samples. The authors also thank KUMC Electron Microscopy Research Lab facility for assistance with the transmission electron microscope.

## Author’s contributions

AC. performed experiments, data analysis and interpretation and preparation of manuscript; KN., XJ., XY., and WXD. performed experiments, data analysis and interpretation; WL, NOD. and WXD. assisted in data interpretation and contributed to the writing of the manuscript; HMN. supervised the project, participated in research design, performed experiments, data analysis and interpretation, and wrote the manuscript. All authors read and approved the final manuscript.

## Abbreviations

AAV: adeno-associated virus
ADRP: adipose differentiation related protein
ALT: alanine aminotransferase
α-SMA: alpha smooth muscle actin
Col1a1: collagen,type-1,alpha-1
CDAHFD: choline-deficient,amino acid-defined HFD (45 % fat) containing 0.1% methionine
COPII: coat protein complex II
CTGF: connective tissue growth factor
ER: endoplasmic reticulum
FFA: free fatty acid
FPLC: Fast protein liquid chromatography
H&E: hematoxylin and eosin
IL-1β: Interleukin-1 beta
IL-10: Interleukin-10
KI: knockin
KO: knockout
LD: lipid droplet
MAM: mitochondrial associated membrane
MTTP: microsomal triglyceride transfer protein
MASLD: metabolic dysfunction-associated steatotic liver disease
MASH: metabolic dysfunction-associated steatohepatitis
PC: phosphatidylcholine
PDI: protein disulfide isomerase
PE: phosphatidylethanolamine
PEMT: Phosphatidylethanolamine N-methyltransferase
PISD: phosphatidylserine decarboxylase
PL: phospholipid
RIPA: radioimmunoprecipitation
SREBP: sterol regulatory element-binding protein
TBG: thyroxine binding globulin
TG: triglyceride
TGF1β: transforming growth factor 1 beta 1
VMP1: vacuole membrane protein 1
WT: wild-type 1

---

**Supplementary Table 1.**
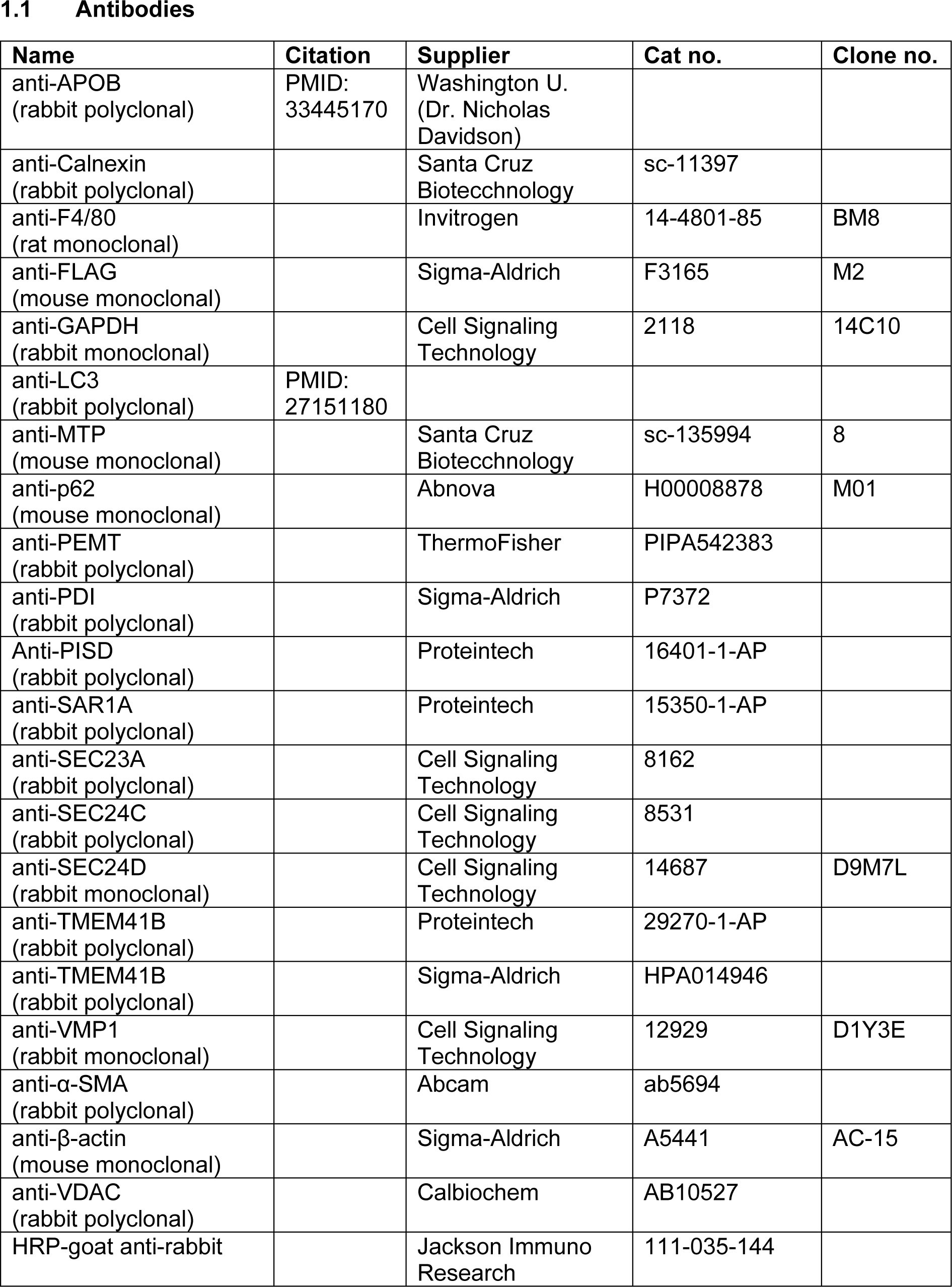
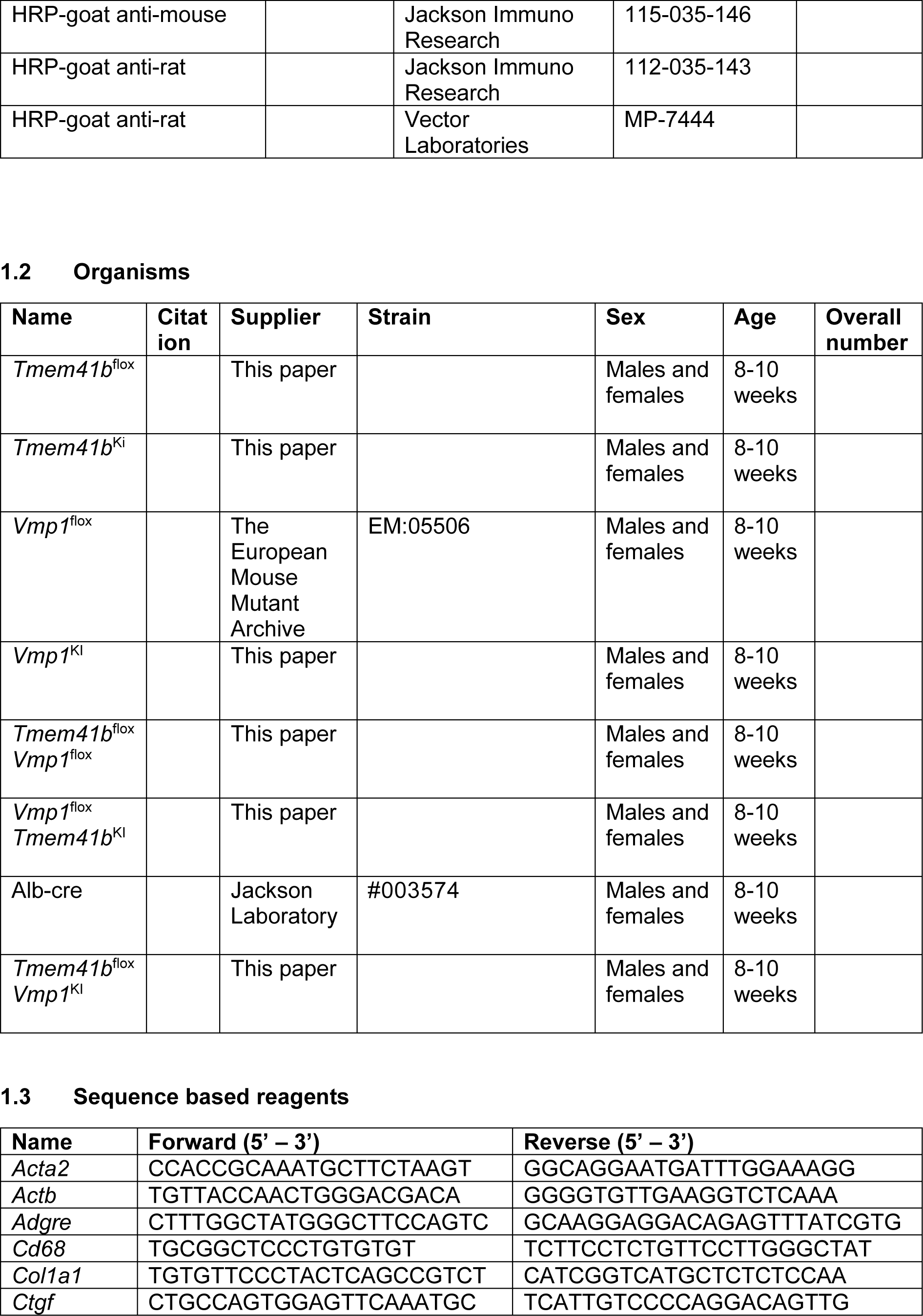
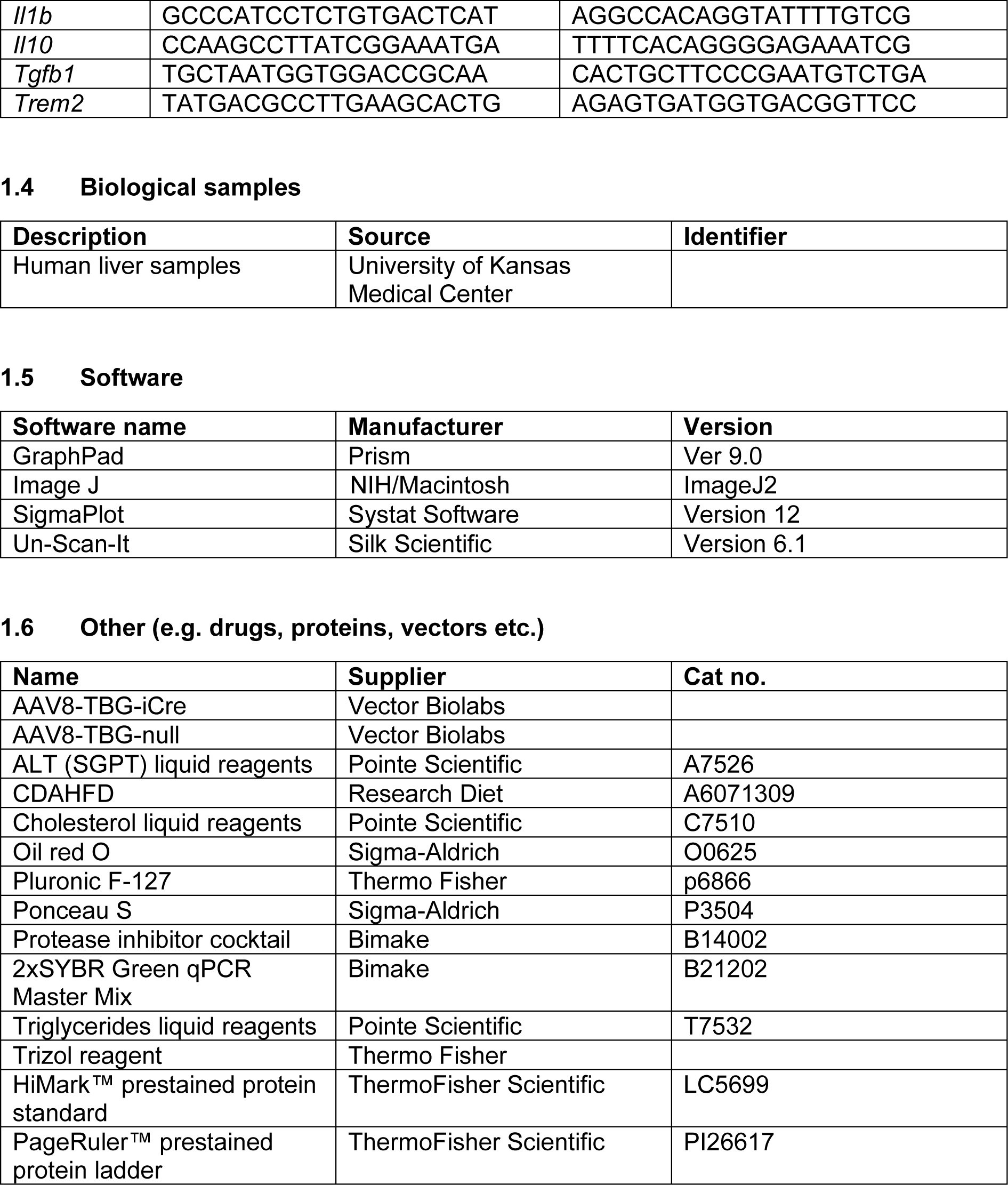
List of antibodies and reagents used in the study.

## Supplementary Figure Legends

**Supplementary Fig. 1.**
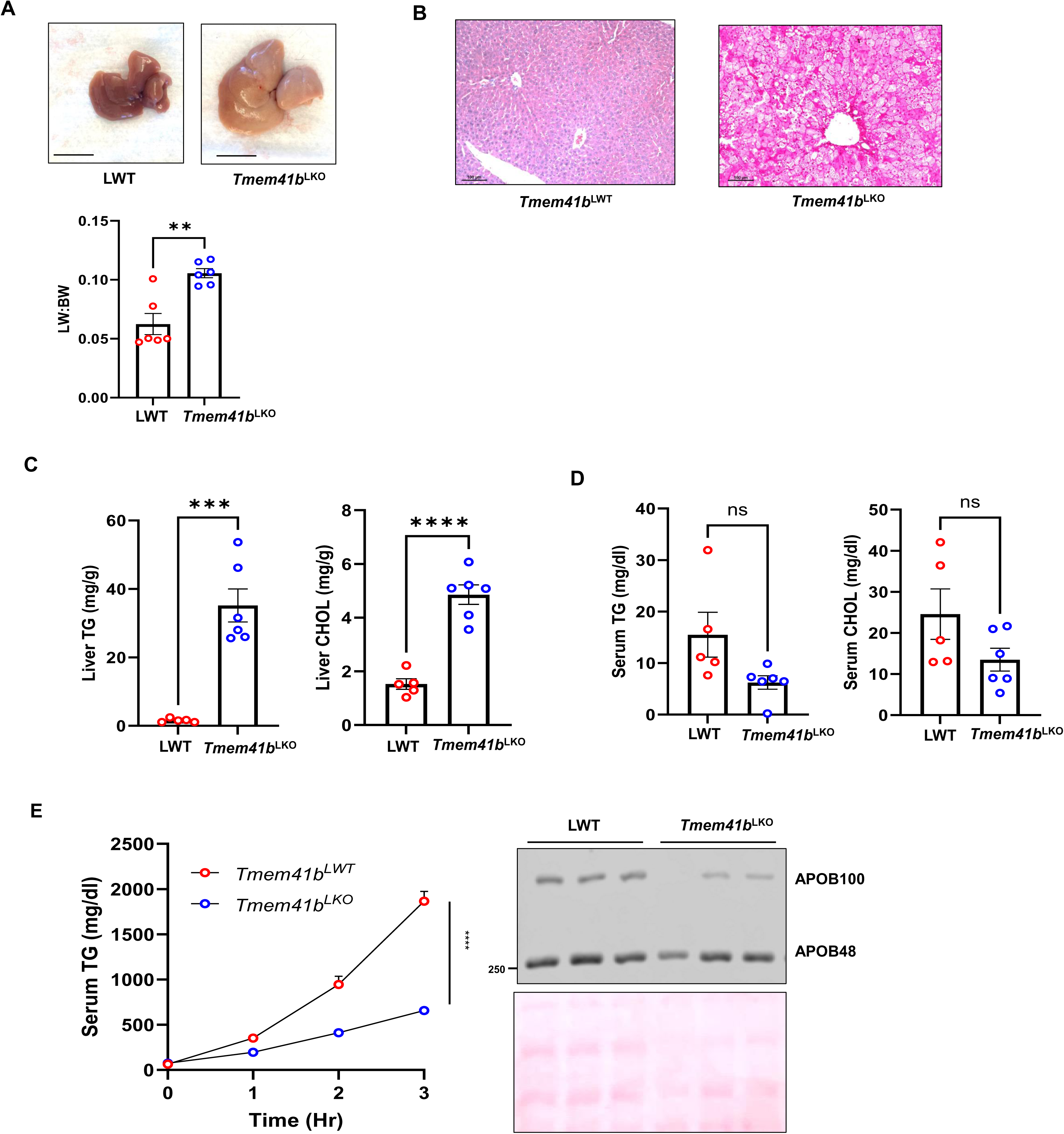
Hepatic deletion of TMEM41B impairs lipoprotein secretion and leads to rapid development of a micro-vesicular steatosis in mice. (A) Representative images of 1-month-old *Tmem41b*^LKO^ and matched WT mice and liver-body weight ratio. Scale bars, 1cm. (B) Representative images of H&E staining of liver tissues from LWT and *Tmem41b^LKO^* mice. Scale bars, 100 µm. Hepatic (C) and serum (D) TG and cholesterol (CHOL) were quantified. (E) LWT and *Tmem41b^LKO^* mice were injected with Pluronic™ F-127 and serum TG concentrations were measured. TG secretion rate bars, 100 µm. Hepatic (C) and serum (D) TG and cholesterol (CHOL) were quantified. (E) LWT and *Tmem41b^LKO^* mice were injected with Pluronic™ F-127 and serum TG concentrations were measured. Serum APOB from LWT and *Tmem41b^LKO^* mice were subjected to immunoblot analysis. Data represent mean ± SEM (n=5-6). Statistical analyses were performed by t-test. ***p<0.001, ****p<0.0001 (unpaired Student’s *t test*).

**Supplementary Fig. 2.**
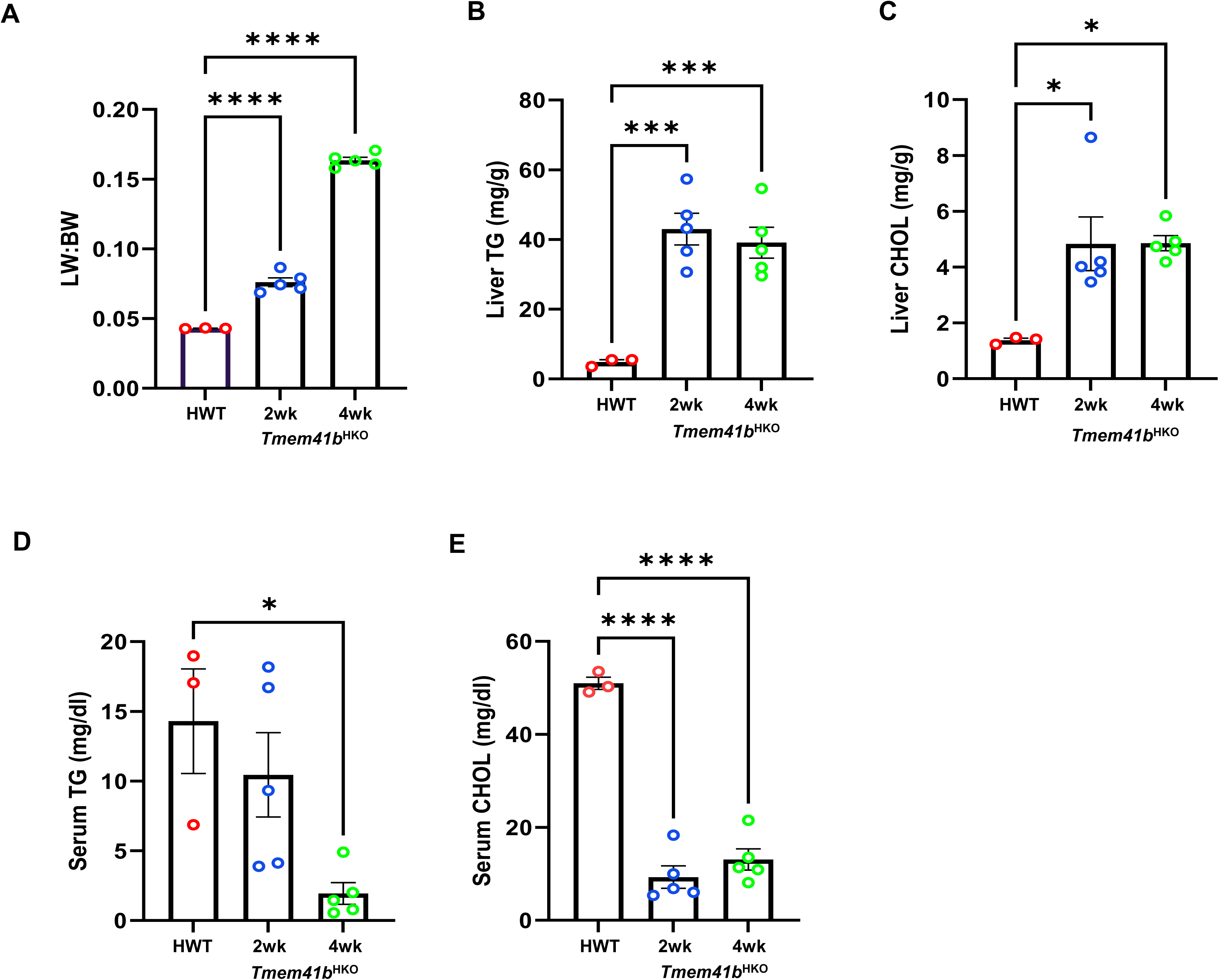
Hepatic deletion of TMEM41B impairs lipoprotein secretion and leads to rapid development of a micro-vesicular steatosis in female mice. (A) Liver/Body Weight ratio in 8-10 weeks old *Tmem41b* female mice at 2 and 4 weeks post AAV8-TBG-cre injection. Hepatic (B-C) and serum (D-E) TG and cholesterol were measured in female mice fed ad libitum of a chow diet. Data represent mean ± SEM (n=3-5). * p<0.05; *** p < 0.001; ****p<0.0001 (one-way ANOVA with post-hoc Turkey test).

**Supplementary Fig. 3.**
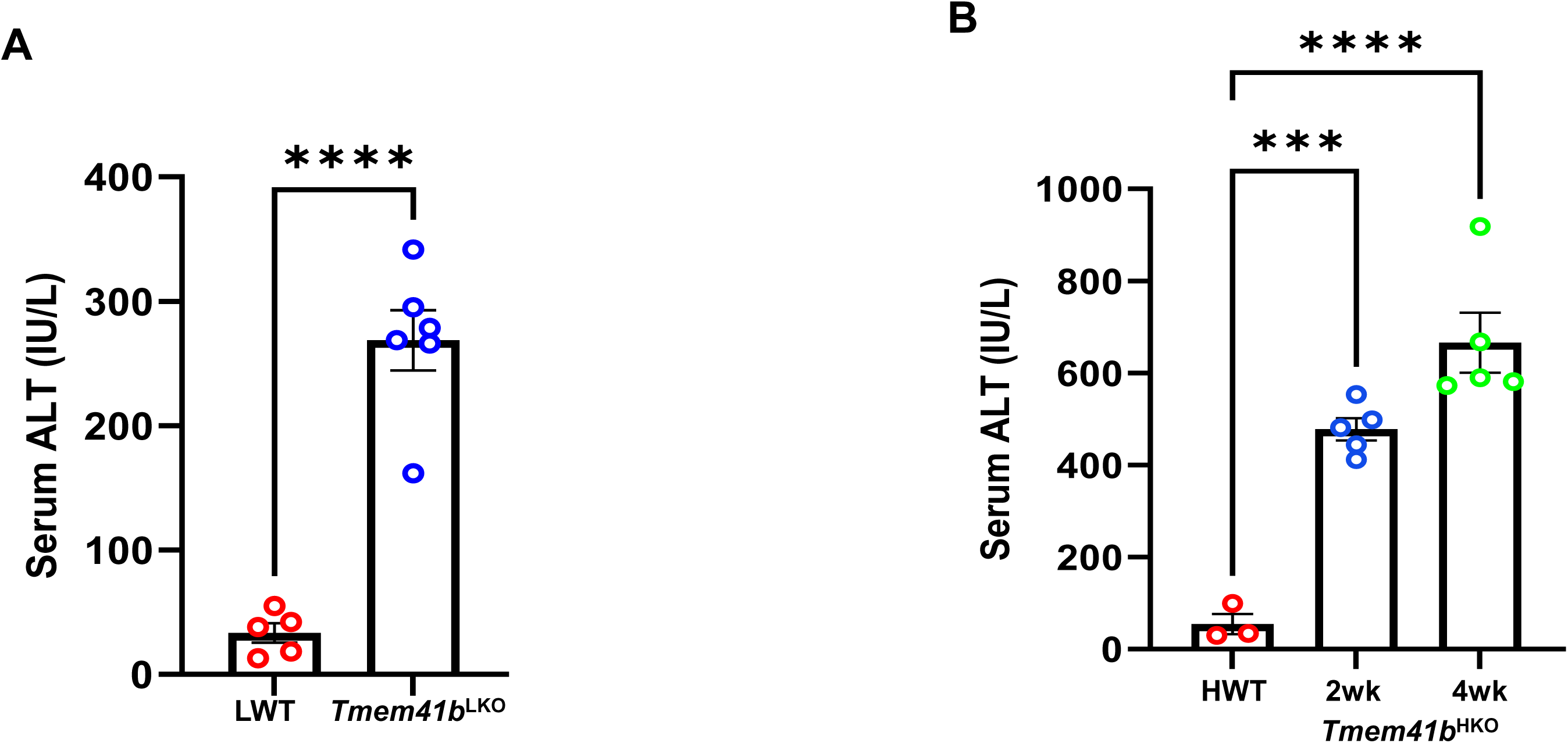
Hepatic deletion of TMEM41B leads to liver injury. (A) Serum ALT activities were measured in 1-month-old *Tmem41b*^LKO^ and matched WT mice (n=5-6). (B) Serum ALT activities were measured in 8-10 weeks old *Tmem41b* female mice at 2 weeks post AAV8-TBG-cre injection (n=3-5). Data represent mean ± SEM. ***p<0.001, ****p<0.0001 (unpaired Student’s *t test* (A) and one-way ANOVA with post-hoc Turkey test (B)).

**Supplementary Fig. 4.**
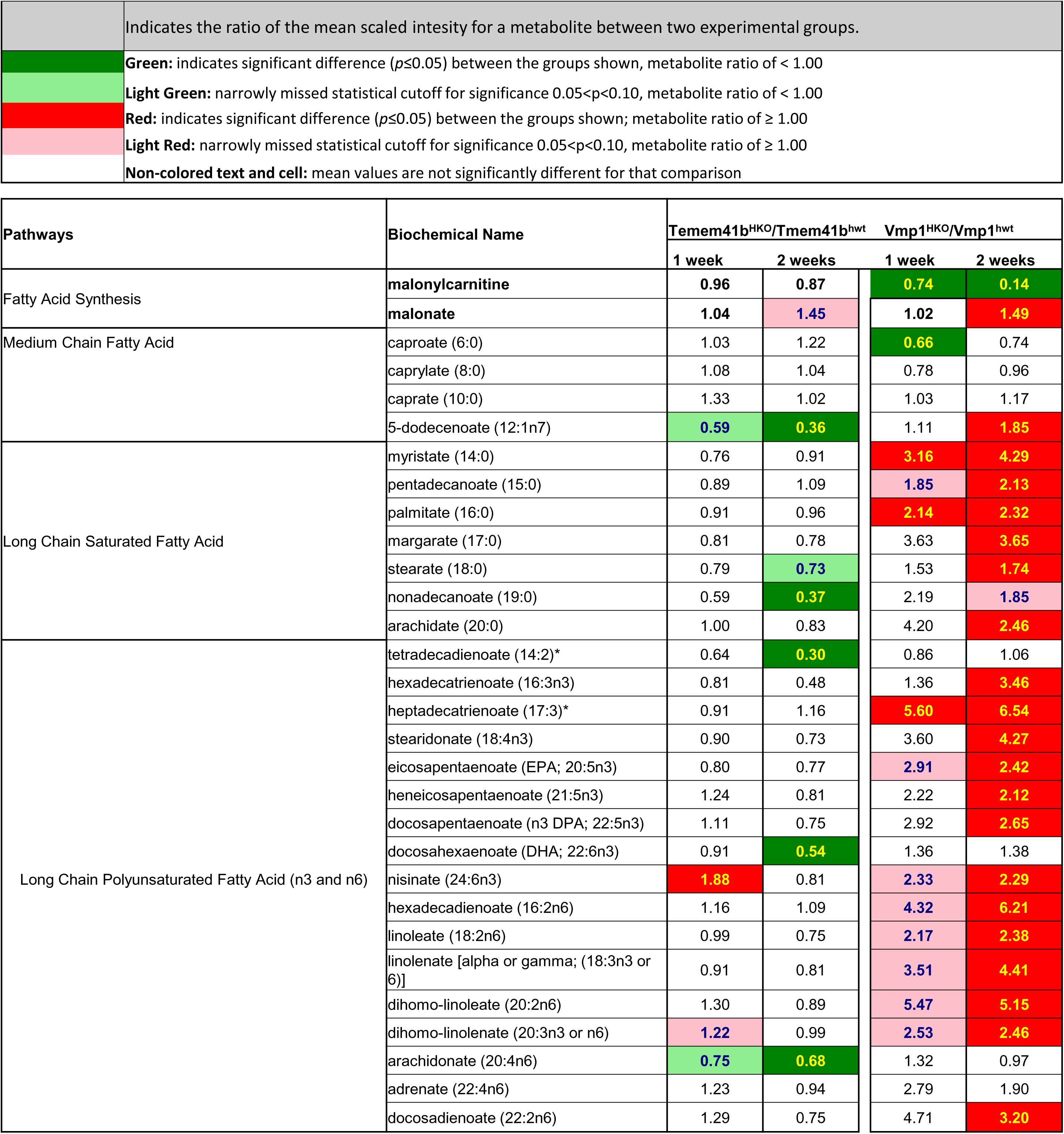
Fatty acid metabolism in hepatic deletion of TMEM41B or VMP1 mouse livers. Heatmap of fatty acids of mouse livers from *Tmem41b* or *Vmp1* mice with 1 and 2 weeks post AAV8-TBG-cre injection by metabolomics analyses.

**Supplementary Fig. 5.**
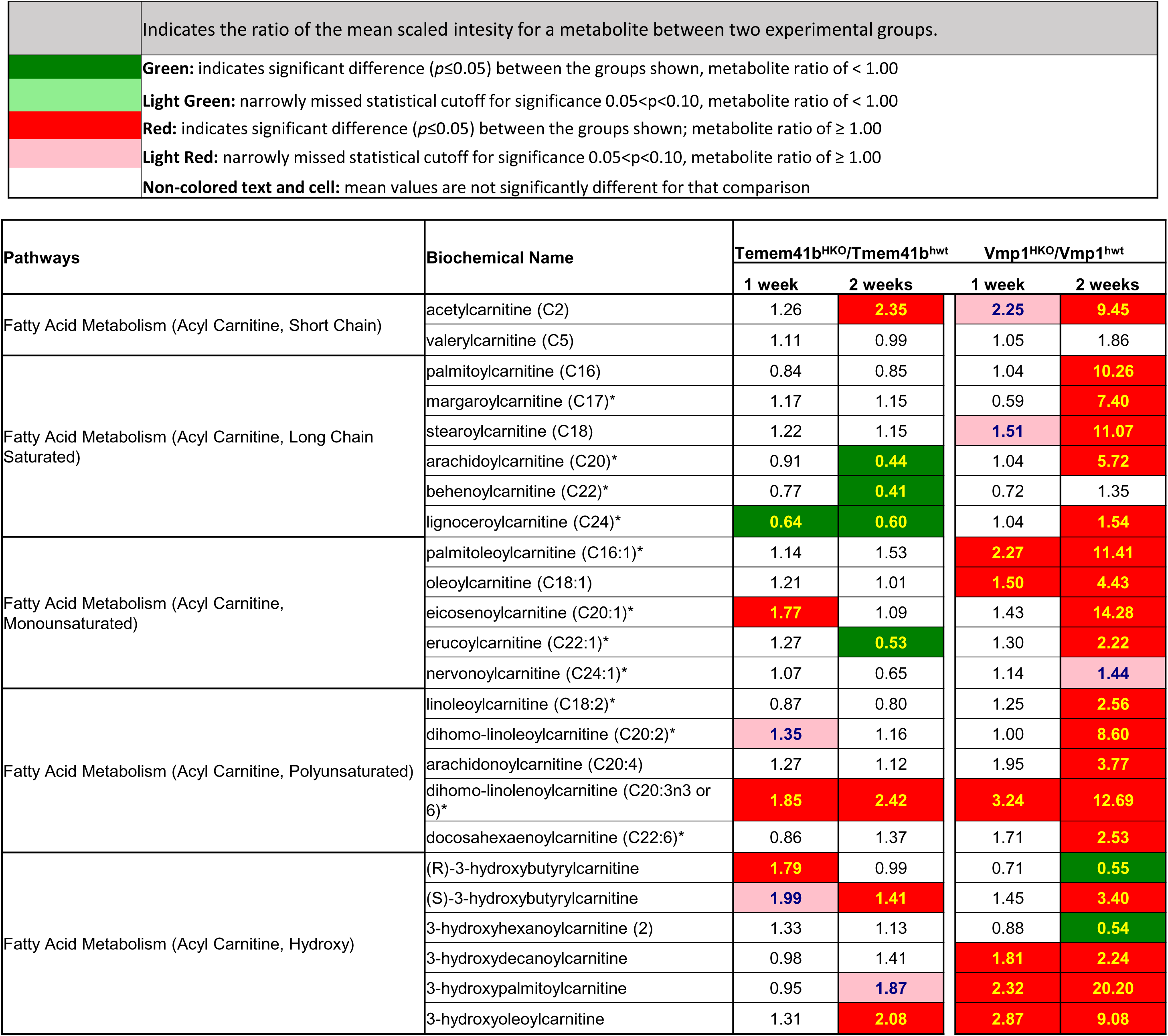
Hepatic deletion of VMP1 increases more acyl carnitine species than hepatic deletion of TMEM41B. Heatmap of acyl carnitine species of mouse livers from *Tmem41b* or *Vmp1* mice with 1 and 2 weeks post AAV8-TBG-cre injection by metabolomics analyses.

**Supplementary Fig. 6.**
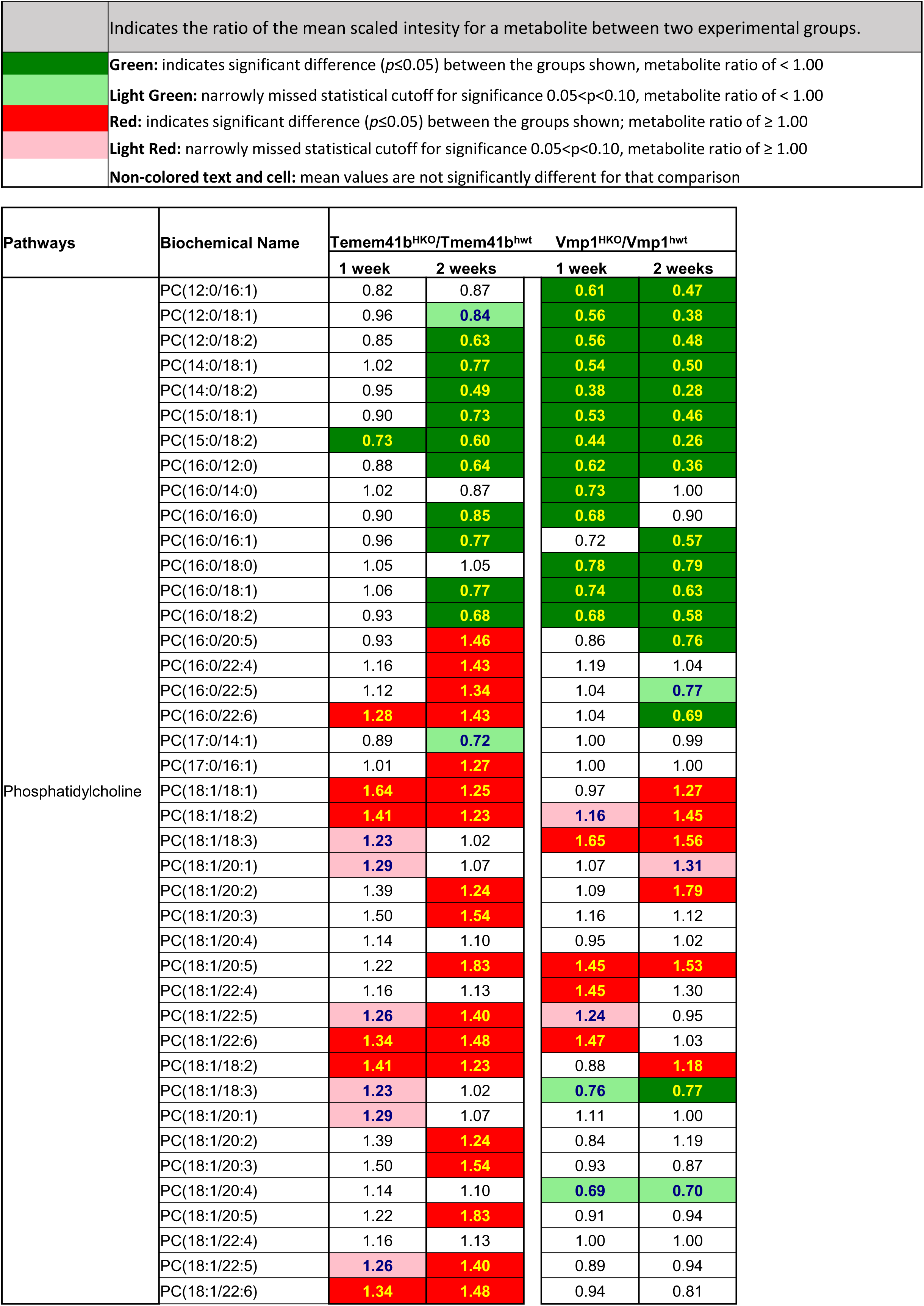
Hepatic deletion of TMEM41B or VMP1 alters PC species in mouse livers. Heatmap of PC of mouse livers from *Tmem41b* or *Vmp1* mice with 1 and 2 weeks post AAV8-TBG-cre injection by lipidomics analyses.

**Supplementary Fig. 7.**
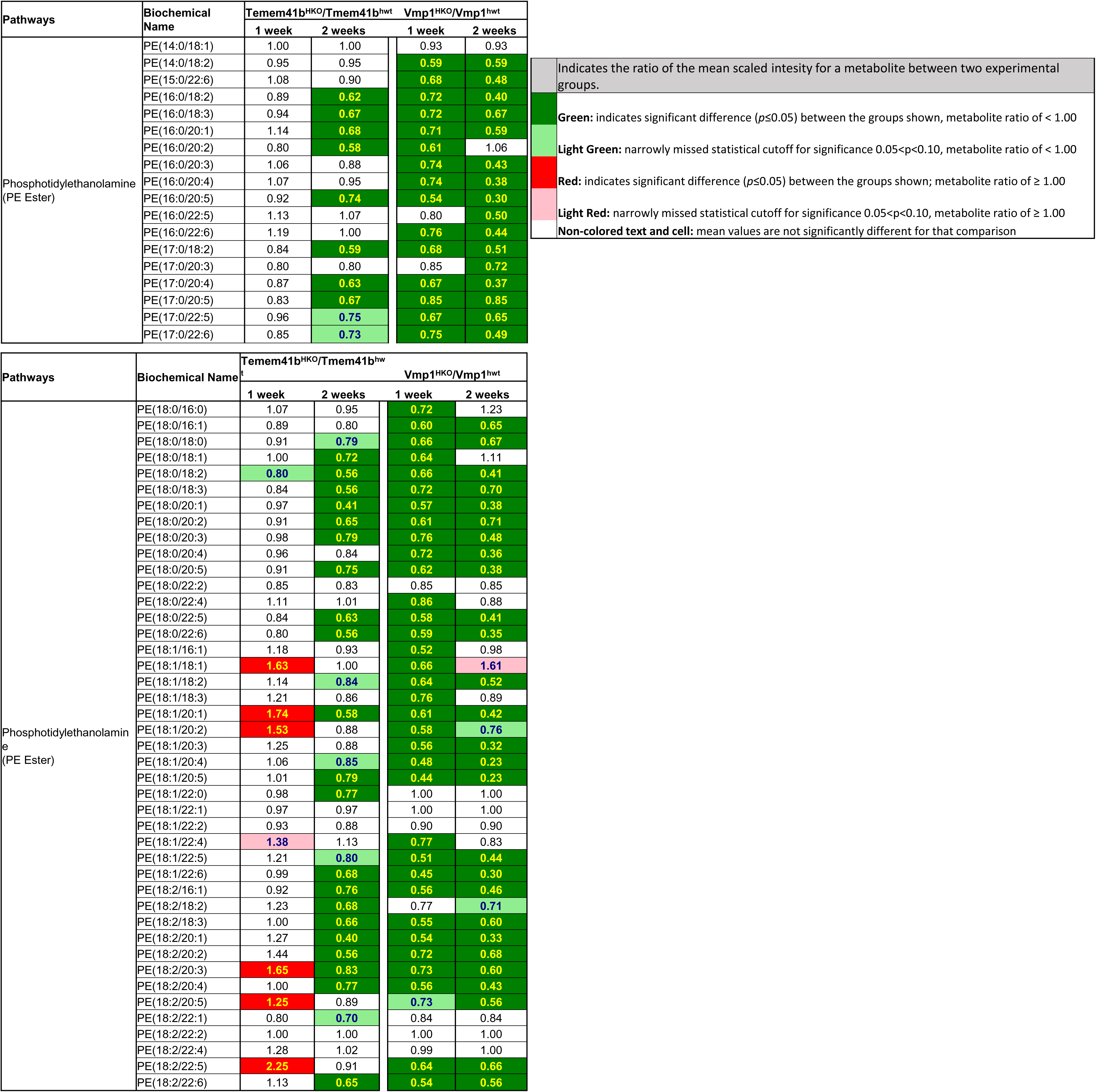
Hepatic deletion of TMEM41B or VMP1 alters PE species in mouse livers. Heatmap of PE species of mouse livers from *Tmem41b* or *Vmp1* mice with 1 and 2 weeks post AAV8-TBG-cre injection by lipidomics analyses.

**Supplementary Fig. 8.**
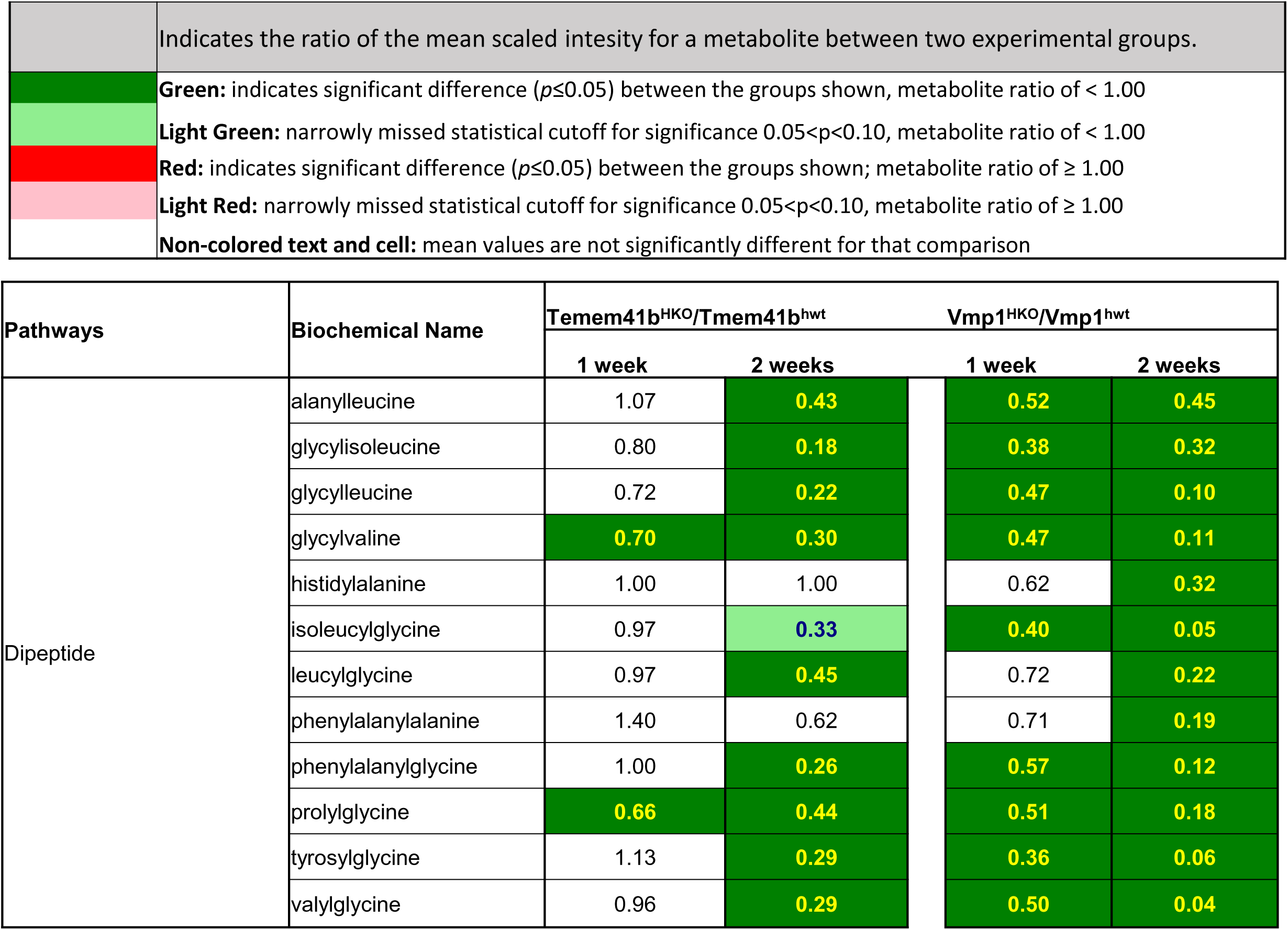
Hepatic deletion of TMEM41B or VMP1 decreases dipeptide in mouse livers. Heatmap of dipeptide of mouse livers from *Tmem41b* or *Vmp1* mice withf 1 and 2 weeks post AAV8-TBG-cre injection by metabolomics analyses.

